# Regulation of the MAPK/ERK system: a computational rule-based model

**DOI:** 10.1101/2025.07.16.665118

**Authors:** Paweł Kocieniewski, Tomasz Lipniacki

## Abstract

MAPK/ERK pathway coordinates multiple cellular functions, including proliferation, apoptosis, and motility, yet it is frequently modeled as a one-purpose cascade. Pathway complexity is a consequence of the existence of isoforms of pathway components that are regulated differently and have specific interacting partners. Here, we propose a rule-based model of the MAPK/ERK pathway that accounts for differential regulation of MEK and RAF isoforms and RAF interactions with 14-3-3 proteins. The model addressed signaling based on the enzymatic cascade as well as regulatory protein-protein interactions. We propose that at low concentrations of growth factors, RAS is activated only in a portion of the membrane. This allows the model to reconcile the observed switch-like and graded responses observed at upper and lower tiers of the pathway, respectively. We demonstrate that functional differences between BRAF and CRAF or ARAF can follow from their different interaction with 14-3-3. The 14-3-3 dimers inhibit all RAF isoforms in close form, but preferentially stabilize BRAF-CRAF dimers to BRAF-BRAF homodimers, and may not stabilize RAF dimers without BRAF. The constructed model allows for the exploration of qualitative differences in MAPK/ERK pathway signaling observed in different cell lines.

## Introduction

The MAPK/ERK pathway (also termed Ras-Raf-MEK-ERK pathway) is a principal channel transmitting signals from growth factors, which typically promote proliferation and survival. This pathway has been intensely studied for the last 40 years due to its fundamental role in cell physiology and involvement in oncogenesis, and as such, its structure, properties, and regulation have been extensively reviewed^1–3^. In brief, growth factors such as EGF or FGF activate growth factor receptors, which dimerize and in turn recruit SOS protein to the cellular membrane. SOS is a guanine exchange factor that activates a small GTPase, RAS, by stimulating the exchange of the RAS-bound GDP to GTP. RAS-GTP subsequently dimerizes and recruits cytosolic RAF kinases (first tier - MAPK), which undergo activation through a series of phosphorylation events and conformational changes that ultimately lead to RAF dimerization and full activation. Active RAF kinases subsequently phosphorylate MEK kinases (second tier - MAP2K), which, in turn, phosphorylate ERK kinases (third tier - MAP3K). The ERK kinases effectuate the physiological response by regulating hundreds of cytoplasmic and nuclear targets through phosphorylation. The ultimate nature of that response is determined by the spatiotemporal pattern of ERK activation as well as the specific set of substrates present in the cell, which depends both on its type and state^4,5^.

While the pathway is generally perceived as linear, with a simple activation flow from one tier to the next, its structure is significantly more complicated. The pathway contains numerous feedback loops. In particular, ERK mediates negative feedback to MEK, RAF, and SOS, enabling transient or oscillatory responses to sustained signals. In particular, this feedback can reset the whole pathway and produce pulsatile responses to transient and tonic signals^6^. This property allows the MAPK/ERK pathway to transmit information with high bit rates^7,8^.

An additional layer of complexity results from the fact that all components of the pathway in vertebrates possess isoforms, i.e., protein variants transcribed from closely related genes (paralogs) originating from a single ancestral gene^9,10^. In this work, we consider two main MEK kinase isoforms (MEK1 and MEK2)^11^ and, in particular, all three RAF isoforms: BRAF, CRAF, and ARAF^12^. The three RAF isoforms share significant sequence homology and overall layout: (1) the N-terminal autoinhibitory domain, which contains the RAS binding domain (RBD) and cysteine-rich domain (CRD) necessary for RAS and membrane recruitment, and (2) the C-terminal domain, which contains the catalytic subdomain and a regulatory N-terminal acidic (NtA) subdomain. Both C- and N-terminal domains contain 14-3-3 binding sites; 14-3-3 proteins are essential regulators of RAF activity, which exist as obligatory homo- and heterodimers of seven known isoforms. Each 14-3-3 has a highly conserved phosphorylation-binding pocket, allowing binding to RAF isoforms’ specific phosphorylated serine residues. Without growth factor signaling, all RAF isoforms exist mainly in a closed, inactive conformation crosslinked by 14-3-3 dimers bound to their C- and N-terminal phosphoresidues. RAFs may bind RAS-GTP dimers only after 14-3-3 dissociates from their N-terminal residues. The RAF dimers formed on RAS platforms may be stabilized by 14-3-3 dimers crosslinking their C-terminal phosphoresidues. This implies that 14-3-3 may stabilize both inactive and active RAF conformations.

The RAF isoforms differ in significant ways. BRAF is most akin to the ancestral RAF kinase; its NtA domain residues, including Ser446, are constitutively phosphorylated, which increases BRAF basal activity and primes it for further activation. In contrast, CRAF and ARAF require recruitment by RAS-GTP for NtA phosphorylation (respectively, at Ser338 and Ser266). Furthermore, unlike BRAF, CRAF can bind and modulate the activity of multiple other proteins through direct protein-protein interactions rather than enzymatic activity. In particular, in its closed conformation, ARAF and CRAF (when phosphorylated at Ser259 and Ser214, respectively) bind and inhibit MST2 kinase, preventing it from participating in the pro-apoptotic Hippo pathway^13^. Engagement of ARAF and CRAF in RAS signaling releases MST2. Thus, growth factors leading to CRAF and ARAF dephosphorylation at Ser259 and SER214 can trigger both proliferation and apoptosis, and the shift of balance towards proliferation is associated with the high activity of AKT, which inhibits MST2^14^ (Romano et al. 2014 - Hippo/CRAF - AKT/ERK). In the “open” conformation, after RAF dimer dissociation, CRAF with phosphorylated Ser338, can form complexes with ROKα, which promotes cytoskeletal rearrangements and motility^15^, and with BAD, Bcl-2, and ASK1 and Bcl-2, inhibiting apoptosis^16–18^.

Regarding regulation by 14-3-3, BRAF recruits 14-3-3 primarily through its high-affinity C-terminal site (Ser729), while lower affinity Ser365 residue on the N-terminus is considered secondary. In turn, the primary 14-3-3 recruitment site of CRAF and ARAF is located at the N-terminus (respectively, at Ser259 and Ser214). Consequently, ARAF and CRAF bind RAS-GTP only after 14-3-3 dissociates from its primary binding N-terminal sites on these isoforms, which likely implies complete dissociation (as the binding to the C-terminal sites, respectively, at Ser621 and Ser582, is unstable). Therefore, neither ARAF nor CRAF may donate 14-3-3 to stabilize RAF dimer. This contrasts with BRAF, which likely binds with RAS-GTP with 14-3-3 bound to its C-terminal domain.

Despite the well-defined architecture, the pathway exhibits significant variability in both behavior and function across different cell lines. From a mechanistic perspective, this can be attributed to order-of-magnitude differences in the levels of protein expression across cell lines. This applies to signal-transmitting cascade components and the ultimate effector proteins, such as ERK-regulated transcription factors. At the systemic level, the characteristic stimulation patterns can differ, implying that ERK activity is either sustained, transient, or oscillatory. Transient activity can be further differentiated based on the duration and steepness of peaks and their regularity or the lack thereof. The significant cell type differences make the universal models with defined parameters unable to reproduce a variety of cell responses.

Due to its importance, the MAPK pathway has been subject to extensive mathematical and computational modeling to understand the role of its structural and regulatory features in performing physiological functions. The first model was published in 1996 by Huang and Ferrel^19^. The authors demonstrated how the three-tier cascade architecture, combined with the dual phosphorylation-based activation of MEK and ERK, produces ultrasensitive behavior, which was later experimentally confirmed in Xenopus oocytes. Over the years, models steadily grew in complexity, incorporating additional features such as feedback loops, scaffold interactions, receptor activation, multisite phosphorylation, and pathway crosstalk.

Feedback regulation is particularly important in shaping the ERK response time profile. The pathway contains numerous negative feedback loops, mostly based on ERK phosphorylation of the upstream pathway components or induction of negative regulators such as phosphatases. Modeling studies have demonstrated their roles in limiting response (transient vs. sustained)^20^ and potentially inducing oscillations^21^. Sturm et al. have also demonstrated that negative feedback can endow the cascade with properties of a negative feedback amplifier, which increases its robustness to internal and external perturbations^22^.

The ERK pathway also contains positive feedback loops, whose functions were predicted to potentiate and sustain ERK activation by weak signals^23^, explain observed switch-like activation of ERK^24^, and potentially introduce bistability. Xiong et al. have indeed experimentally verified that a positive feedback loop (ERK-Cdc2) underlies Xenopus oocyte maturation and concomitant all-or-none activation of ERK^25^; they have also demonstrated the system to be bistable due to the combination of this positive feedback loop with the non-linear activation of ERK, providing a basis for irreversible cell fate decisions and memory. Another positive feedback loop is based on the upregulation of SOS GEF activity due to the allosteric binding of RAS-GTP^26^ . This feedback was theoretically demonstrated to drive, in the case of slow diffusion, membrane clustering and activation of traveling waves^27^ ; it was experimentally shown to drive a digital, hysteretic RAS activation in lymphoid cells^28^.

Computational studies also uncovered non-obvious sources of positive feedback that do not require explicit upregulation but emerge from substrate-based enzyme saturation or sequestration. In particular, Markevitch et al. demonstrated an effective positive feedback based on the dual (de)phosphorylation cycle of the enzymes in the cascade, which can yield bistability^29^. They then demonstrated how this mechanism could underlie long-range propagation of ERK activity in the cell^30^. Legewie et al. later proposed another model with positive feedback stemming from enzyme-substrate interactions, where fully phosphorylated ERK dissociates from MEK and increases its fraction^31^ available for activation by RAF.

Combinations of negative and positive feedback potentially produce even more complex behavior. Yasemi et al. showed how the SOS-RAS positive feedback introduces bistability and switching dynamics, while the addition of negative feedback from ERK to SOS introduces sustained and damped oscillations. Kochańczyk et al. 2017 have demonstrated how a fast SOS-RAS positive feedback loop embedded in a slow negative feedback loop (ERK to SOS) can produce relaxation oscillations and enable frequency rather than amplitude-based encoding^32^. Marrone et al. have shown more generally how positive and negative feedback jointly ensure either (1) relaxation-type oscillations or (2) smoother oscillations in a narrow range of frequencies^33^.

Few models have addressed the role of isoforms in the cascade. Harrington et al. demonstrated how different trafficking constants between active ERK1 and ERK2 can affect their intracellular distribution (cytoplasm vs. nucleus) and, therefore, cell response^34^. In another early model, Robubi et al. incorporated the then-known differences between BRAF and CRAF in their (in)activation rates and enzymatic activities; the model suggested that both isoforms equally affect the amplitude, but BRAF is more responsible for the response duration^35^. Subsequently, Kocieniewski et al. accounted for the differences in negative feedback regulation of MEK1 and MEK2. They demonstrated that the amplitude of ERK response can be controlled by the total amount of MEK1 and MEK2 while their ratio controls response duration^36^.

In this study, following our earlier models (Kochańczyk et al. 2017 and Varga et al. 2018), we construct a rule-based MAPK system model that captures the combinatorial complexity of RAF isoforms signaling of the MAPK/ERK system^32,43^. We exploit the model to investigate (1) switch-like and gradual activation of respectively lower (MEK and ERK) and upper tiers (RAS and RAF) of the signaling cascade, (2) regulation of CRAF competence to interact with ROKα, and MST2, and (3) differential regulation of BRAF, CRAF, and ARAF isoforms by 14-3-3 proteins.

## Results

### Rule-based model of MAPK pathway

The proposed model expands the MAPK pathway model constructed by Kochańczyk et al. 2017 and later developed by Varga et al. 2017 to elucidate the dynamics of CRAF-ROKα interactions^32,43^. Following the original model, the new model accounts for one positive feedback from RAS to SOS that is responsible for bistability (observed in the absence of negative feedbacks), and three negative feedbacks mediated by ERK and involving MEK1 (and indirectly MEK2), RAFs, and SOS. The last feedback loop from ERK to SOS is of key importance as it allows for relaxation oscillations, while the feedbacks to MEK and RAFs mainly modulate the shape of these oscillations.

The model is focused on the specificity of MEK and (mainly) RAF isoforms regulation and their complex interaction with 14-3-3 proteins. To account for these interactions, we employed rule-based modeling^37,38^ that allows for defining systems with a large number of species (for example, phosphostates of proteins and their complexes) and are regulated by a large number of biochemical interactions using a smaller number of rules. In the study, we do not aim to fit parameters to reproduce the behavior of cells of a particular type but instead use the proposed model as a platform to investigate qualitative changes in cell responses that may result from changes in particular parameters.

The key interactions of MAPK pathway components are visualized in Fig. 1. The BioNetGen specification of the computational model, together with the auxiliary numerical codes used for the model analysis, are provided as Supplementary Code S1. The species specification and default parameters are summarized in Supplementary Text S1. The details of the model structure and numerical simulations are provided in Materials and Methods.

**Figure 1.**
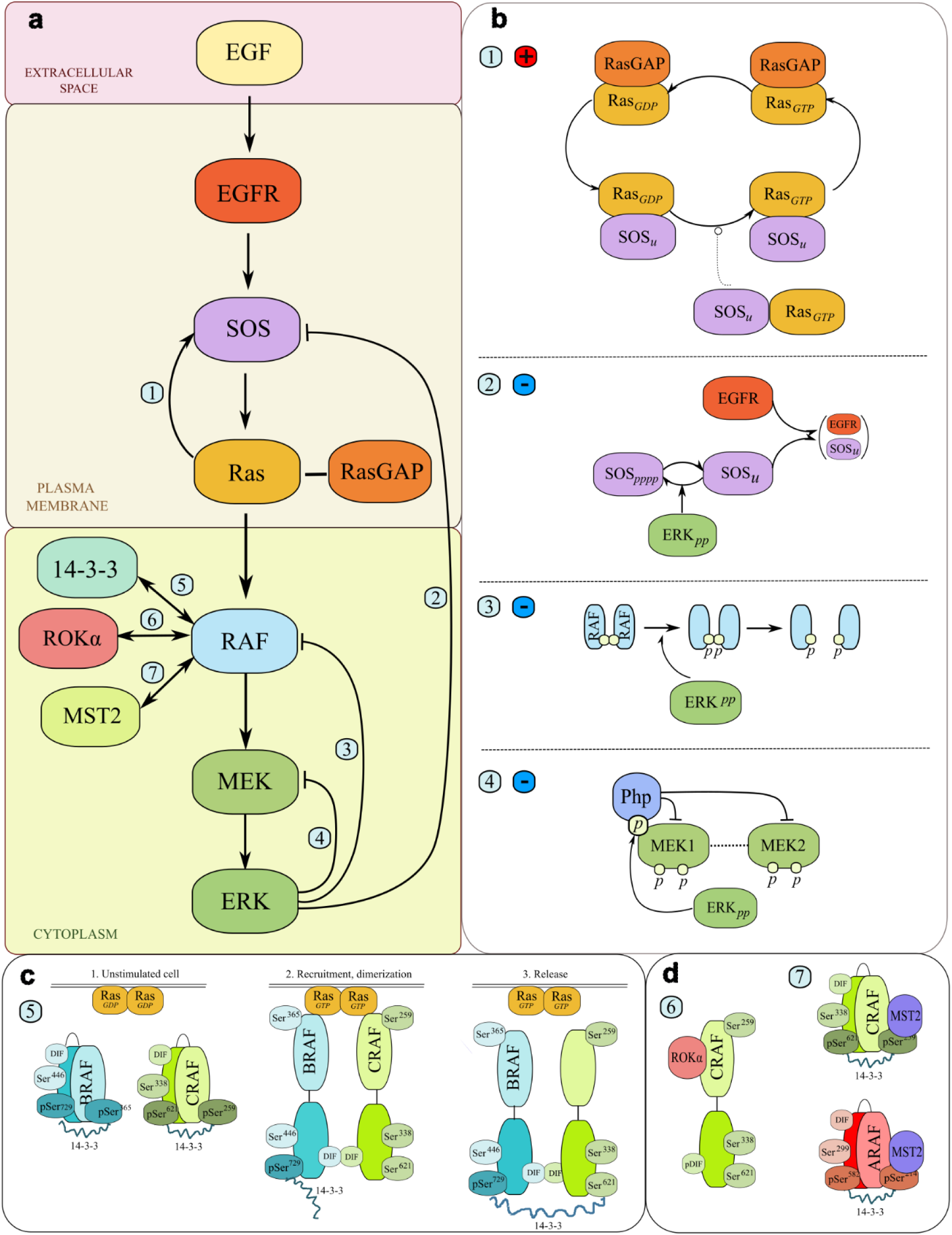
Scheme of the model. **(a)** The diagram of the RAF/ERK pathway. The model accounts for MAPK/ERK pathway signaling from the level of extracellular EGF stimulation through the distinct membrane and cytoplasmic compartments. It comprises all the tiers of the cascade down to ERK and includes three RAF isoforms (BRAF/CRAF/ARAF) as well as two isoforms of MEK (MEK1/MEK2). **(b)** The feedback loop architecture. The model accounts for one positive feedback loop (1) SOS-RAS and three negative feedback loops from ERKpp to (2) SOS, (3) RAF, and (4) MEK. **(c)** The RAF kinases protein-protein interactions. The model accounts for interactions of the RAF isoforms with 14-3-3 proteins. In particular, BRAF initially binds 14-3-3 via its C-terminus, while CRAF and ARAF through their N-terminus. In quiescent cells, all RAF kinases exist in the “closed” conformation, crosslinked by 14-3-3. In stimulated cells, RAF kinases in the “open” conformation without 14-3-3 bound to their N-terminus are recruited by RAS-GTP platforms and form homo- and heterodimers. Since BRAF can retain 14-3-3 at its C-terminus, 14-3-3 can crosslink BRAF with CRAF, ARAF, or other BRAF, stabilizing B-B, B-C, and B-A dimers. In contrast, C-C, A-A, and C-A homo- and heterodimers cannot be stabilized via 14-3-3 crosslinking. **(d)** The CRAF and ARAF protein-protein interactions. When phosphorylated at their N-terminal 14- 3-3 binding sites (Ser259 and Ser214), both CRAF and ARAF can bind and sequester MST2 kinase (6). When CRAF is in its open conformation and phosphorylated on the NtA domain (Ser338), it can bind and inhibit ROKα and ASK1 (7); feedback phosphorylation at the dimerization interface (DIF) by ERKpp disrupts dimers and enhances these interactions.

### Bistability and relaxation oscillations

As in the Kochańczyk et al. 2017 model^32^, system bistability, observed in the absence of negative feedbacks, is a consequence of positive feedback coupling SOS and RAS together with nonlinearity in RAS-GTP to RAS-GDP conversion associated with low abundance of ‘converting enzyme’ RAS-GAP. Such architecture results in switch-like responses of upper tiers of the signaling cascade: RAS and RAF, meaning abrupt growth of activity of these components from the low to the high levels (Fig. 2a). Experimental data indicate, however, that although ERK activation has typically a switch-like character^24,39,40^, the activity of RAS and RAF (when observed at whole cell level) grows more gradually with signal strength^14,15^. To reconcile these observations, we propose that at low EGF concentration, switch-like activation of RAS occurs only on a fraction of the cellular membrane, *f*, growing with EGF concentration. We chose equal *f* = *EGF*/(*EGF +* M_EGF_) with M_EGF_ = 300 pg/ml; the M_EGF_ value defines the EGF range for which gradual growth RAS and RAFs activity is observed, which can vary between cell lines. Such localized RAS activations are observed experimentally in response to various stimuli, and enable RAS to coordinate cytoskeletal reorganization, cell-cell adhesion, and eventually cell motility^41,42^. As the bistable RAS switch occurs only on a portion of the membrane, at the whole cell level it produces just a small jump of RAS activity (Fig. 2b). However, because of signal amplification at the RAF, MEK, and ERK levels, activation of a relatively small fraction of RAS leads to nearly full activation of MEK and ERK; when RAS switches to its upper branch, the doubly phosphorylated MEK and ERK (MEKpp and ERKpp) reach more than 90% of whole protein levels. Consequently, we observe nearly gradual activation of RAS and RAF and switch-like activation of MEK and ERK (Fig. 2b). Because of a high signal amplification downstream of RAS, its oncogenic mutations leading to the constitutive activation are difficult to counteract at lower tiers of the pathway.

**Figure 2.**
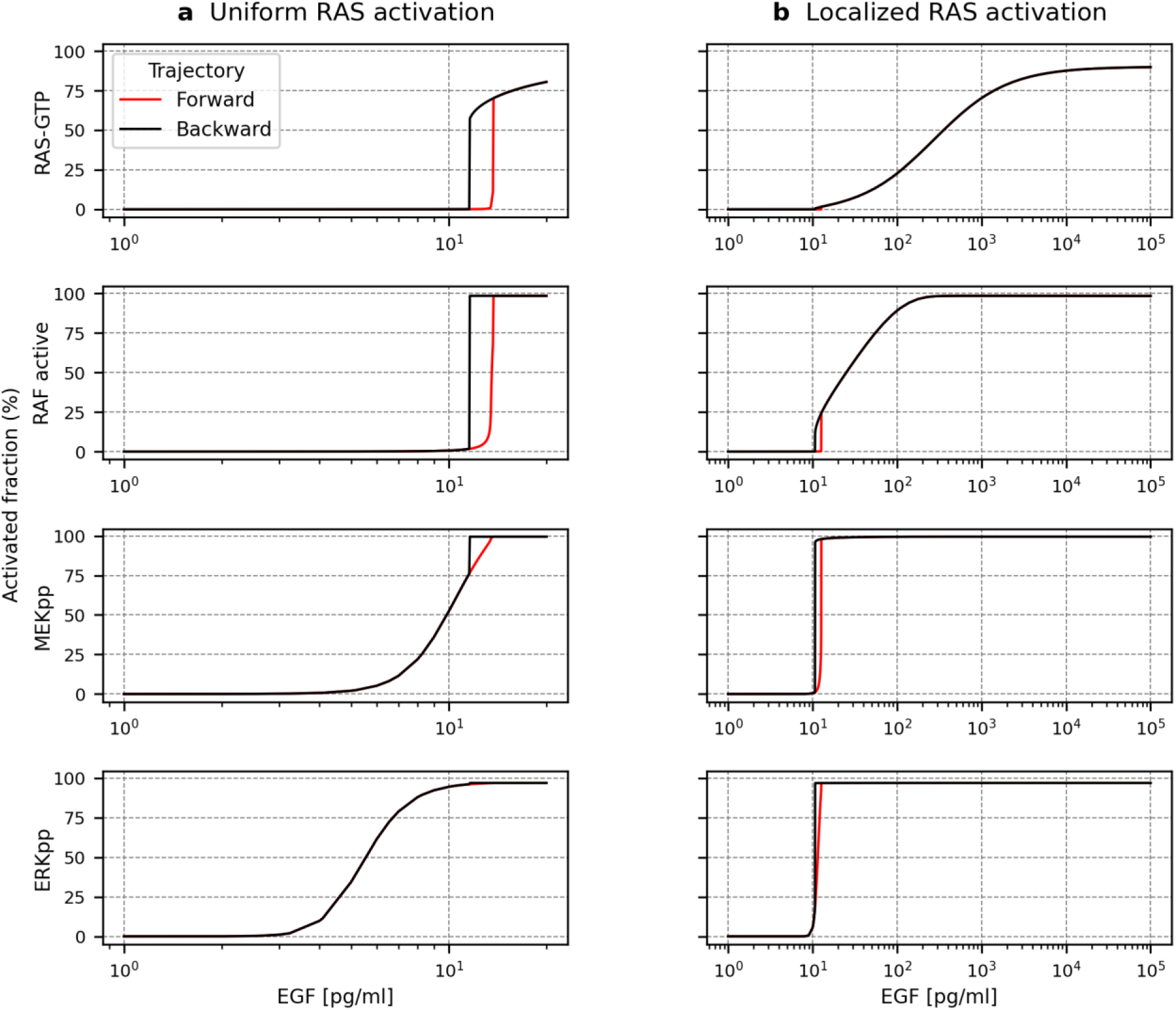
Bistable switching with all negative feedback loops removed. Forward and backward activation profiles of RAS-GTP, active RAFs, MEKpp, and ERKpp in response to, respectively, increasing and decreasing concentration of EGF. Active RAF stands for all dimerized RAF isoforms molecules that are phosphorylated at both their NtA domains and C-terminal 14-3-3 binding sites. **(a)** For uniform RAS activation abrupt switch is observed for RAS-GTP and active RAF **(b)** For RAS activation restricted to a portion of the membrane (growing with EGF concentration), an abrupt switch is observed for MEKpp and ERKpp. Notice the different horizontal scales in the two panels.

In both model variants, the bistability region is relatively narrow and extends for EGF concentrations between approximately 10 and 12 pg/ml. From now on, we focus on the (nominal) model in which RAS activation is localized to the portion of the cell membrane. The negative feedback from ERK to SOS encompasses positive feedback from SOS to RAS and frustrates bistability to produce relaxation oscillations, Fig. 3a. As expected for such systems, oscillations are born in saddle-node on invariant circle (SNIC) bifurcation with an infinite period^32^. For the nominal model parameters, limit cycle oscillations exist for a range of EGF concentrations: (from ∼12 to ∼1500 pg/ml). As shown in Fig. 3b the period is the lowest for intermediate EGF concentrations and then increases with the strength of stimulation. For EGF concentration larger than ∼1500 pg/ml, oscillations are damped (the profile of such damped oscillations is shown in Fig. 3a for EGF concentration equal 10^4^ pg/ml. The range of EGF concentrations for which limit cycle oscillations are observed depends on the strength of negative feedback from ERK to SOS, and in certain range of the negative feedback strength, the limit cycle oscillations are observed for arbitrarily large EGF concentrations. This implies that cells of a particular type can be tuned to respond in a pulsatile manner to EGF stimulation up to some arbitrary concentration. As the temporal response patterns dictate physiological outcomes (like proliferation or differentiation^40^), the same stimuli can lead to different outcomes depending on cell type^5^.

**Figure 3.**
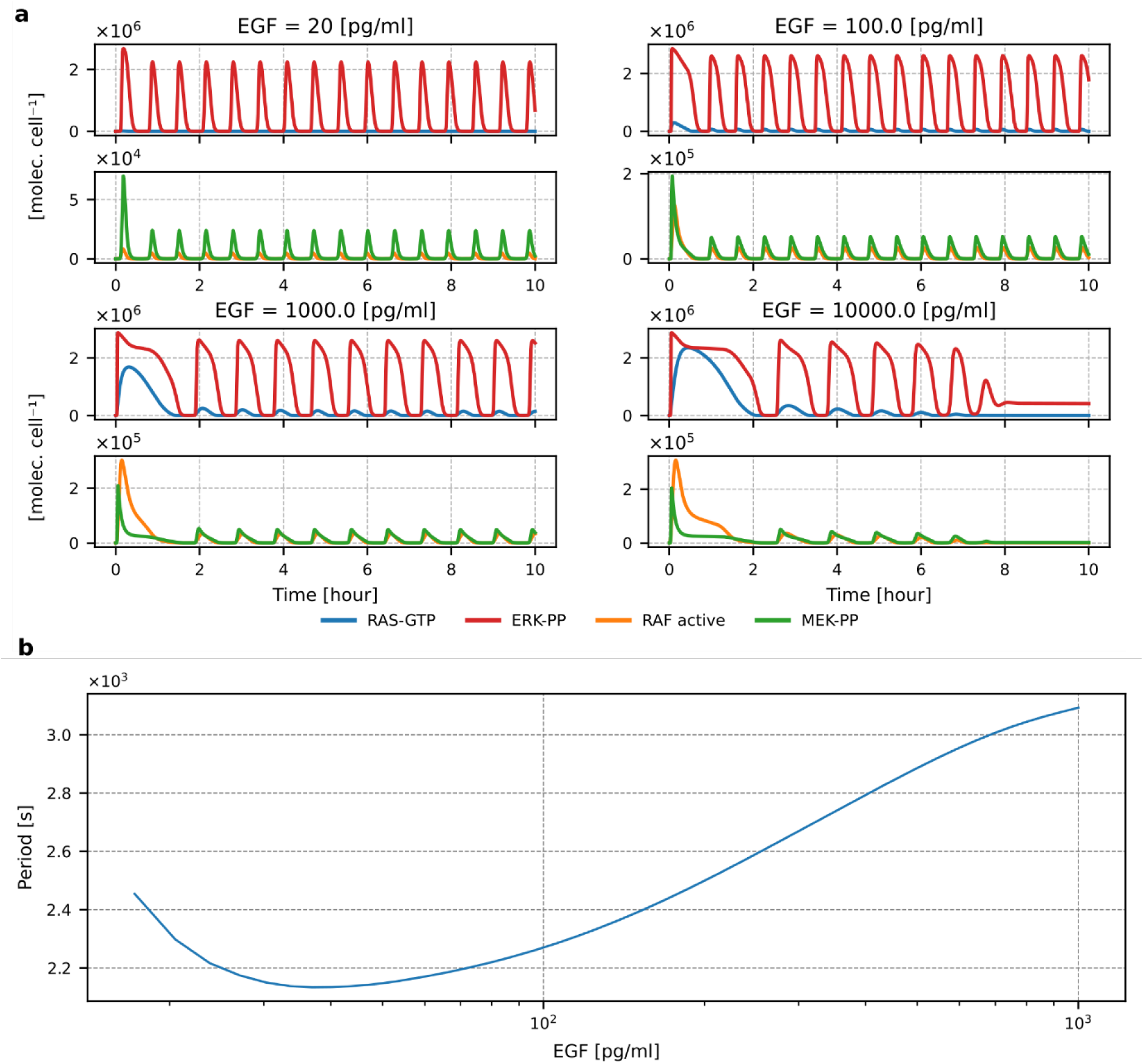
Relaxation oscillations of the MAPK system. **(a)** Relaxation oscillations profiles of RAS-GTP, active RAF, MEKpp, and ERKpp for four EGF concentrations as indicated. Notice the different vertical scales on the subpanels. **(b)** Oscillation period as a function of EGF concentration.

The dependence of the first and subsequent peak amplitudes on EGF concentration differs for the four MAPK/ERK cascade tiers. The amplitude of ERKpp oscillations remains nearly independent of EGF concentration, and the first peak is only somewhat higher than subsequent ones. Nearly all-or-nothing ERK activation is crucial for whole system regulation, as it allows ERK to terminate both weak and strong signals. The MEKpp first peak amplitude remains nearly unchanged for EGF concentrations higher than 100pg/ml and is higher than the amplitudes of subsequent peaks. For RAS and RAF, the first peak amplitude grows with EGF concentration, reaching 25% and 60% of activated protein, respectively, and is higher than the amplitude of subsequent peaks. This allows both RAS and RAF to play additional, dose-dependent roles beyond transmitting signals to MEK and ERK. With this respect, RAS coordinates both ERK and AKT activity as well as cell motility; CRAF, depending on its phosphorylation status, may form complexes with ROKα and ASK1, regulating cell motility and inhibiting apoptosis, and both ARAF and CRAF form complexes with MST2, regulating apoptosis^13,14,43^.

### Activation of RAS, ERK proteins, RAF, and MEK isoforms

The existence of three RAF and two MEK isoforms implies that the signal from RAS to ERK has several paths to follow. Although only MEK1 possesses the ERK phosphorylation site (Thr292) and mediates negative feedback from ERK, the activation profiles of MEK1 and MEK2 are nearly identical (Fig. 4 and Supplementary Figure S1). This is because ERK- phosphorylated MEK1 may bind the MEK-specific phosphatase that dephosphorylates MEK1 as well as MEK2 when the latter is in a heterodimer with MEK1.

**Figure 4.**
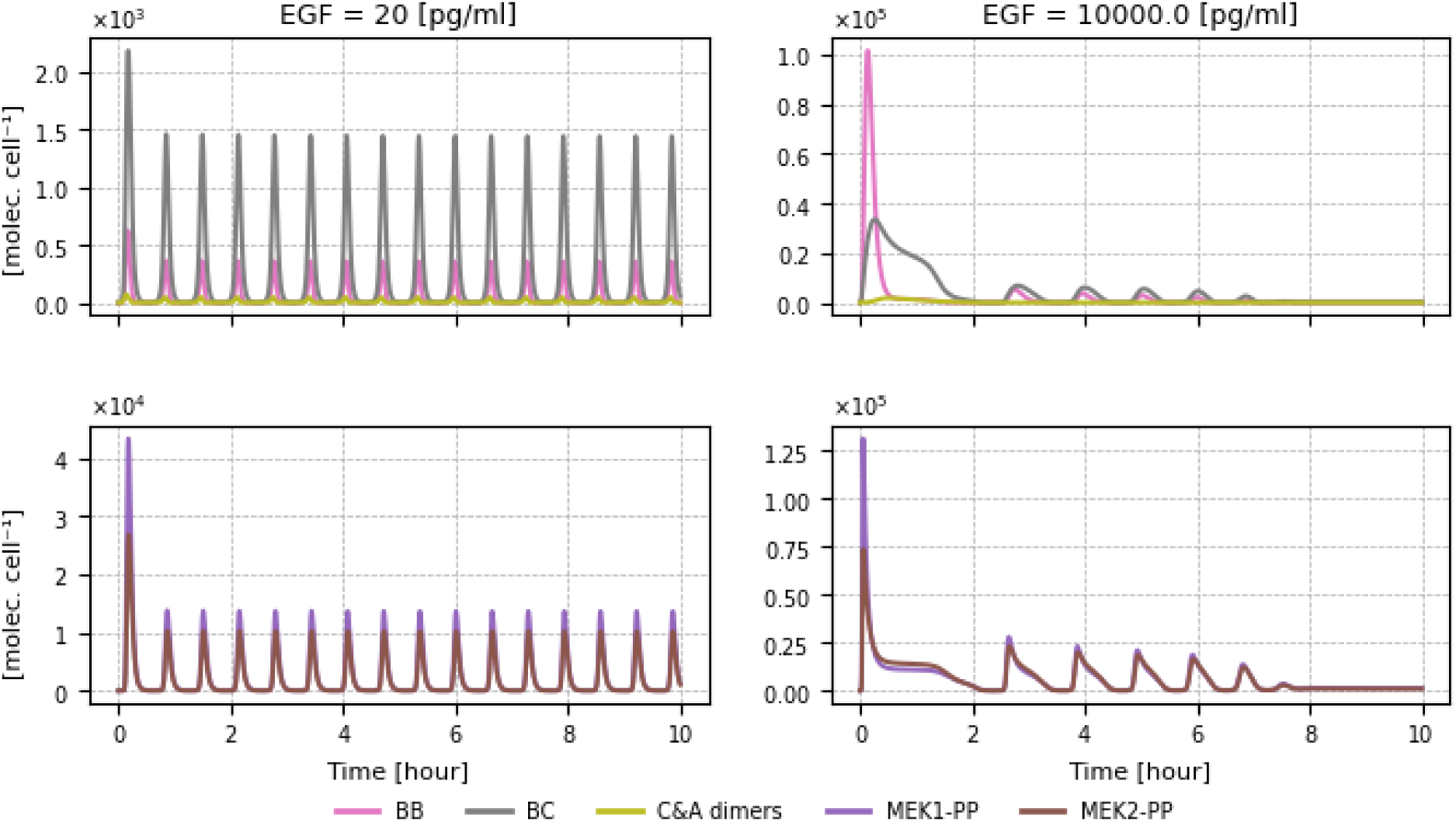
Relaxation oscillations profiles of RAF and MEK isoforms. The plot shows abundances of BB, BC, and C&A RAF dimers, and MEK1pp, MEK2pp molecules for low and high EGF concentrations. C&A dimers stand for all RAF dimers not containing BRAF. The corresponding plots for the intermediate EGF concentrations, 100 pg/ml and 1000 pg/ml, are provided in Supplementary Figure S1.

In turn, RAF isoforms, which may form both homodimers and heterodimers, are regulated distinctively (Fig. 4). For the sake of simplicity, we have assumed the same rules and constants for ARAF and CRAF regulation, but different rules for BRAF. The difference between BRAF and the two other isoforms is in their interaction with 14-3-3 dimers. To recapitulate, in the absence of a signal, all three isoforms are mainly in their closed forms, being crosslinked by 14-3-3 dimers, which are bound to their (phosphorylated) N and C-terminal phosphosites. In turn, when activated, RAF dimers may be stabilized by 14-3-3 dimers bound to RAFs’ C-terminal phosphosites.

The difference between RAF isoforms follows from the fact that the primary binding site for 14-3-3 on BRAF (S729) is on its C-terminal, while the residue on the N-terminus (S365) is considered secondary. In contrast, for ARAF and CRAF, the primary 14-3-3 binding sites (respectively S214 and S259) are on the N-terminus, while secondary binding sites (S582 and S629) are on the N-terminus^43–45^. To reflect the lower affinity of binding to secondary sites, it is assumed (in the model) that when 14-3-3 binds RAF monomer, it must first bind the stronger primary site, then the secondary, and when it dissociates, it first dissociates from the secondary site. Since all three RAF isoforms bind RAS using their N-terminal sites, ARAF and CRAF may bind RAS only when they are not in complex with 14-3-3 (because 14-3-3 binding interferes with RAS and membrane recruitment). In turn, BRAF can bind RAS, having a 14-3- 3 dimer bound to its C-terminal sites. This has pronounced dynamical consequences: The 14- 3-3 dimer associated with BRAF bound to RAS can then bind the ARAF, CRAF, or other BRAF stabilizing BRAF-ARAF or BRAF-CRAF heterodimers or BRAF-BRAF homodimers (provided that the C-terminal site on the second BRAF is free). This difference implies that BRAF-ARAF, BRAF-CRAF, or BRAF-BRAF dimers can be stabilized by 14-3-3, and therefore, as shown in Fig. 4, are much more abundant than dimers that do not contain BRAF (i.e., CRAF-CRAF, ARAF-ARAF, and CRAF-ARAF, jointly termed C&A dimers).

Although BRAF-BRAF homodimers as well as BRAF-ARAF and BRAF-CRAF heterodimers may be stabilized by 14-3-3, their dynamics are different (Fig. 4 and Supplementary Figure S1). Heterodimers are formed more eagerly at low RAS-GTP levels (i.e. at low EGF concentration or during recurrent RAS activity pulses), and are more stable, which also follows from the assumed interactions with 14-3-3. The different activity time profiles observed for BRAF-BRAF monomers and BRAF-CRAF (or BRAF-ARAF) heterodimers observed for 10^4^ pg/ml EGF concentration likely lead to dampened oscillations. In cells deficient of CRAF and ARAF, in which the whole signal to ERK is transmitted by BRAF-BRAF monomers, the limit cycle oscillations are observed for 10^4^ pg/ml EGF (Fig. 5a), and continue for arbitrary large concentrations of EGF.

**Figure 5.**
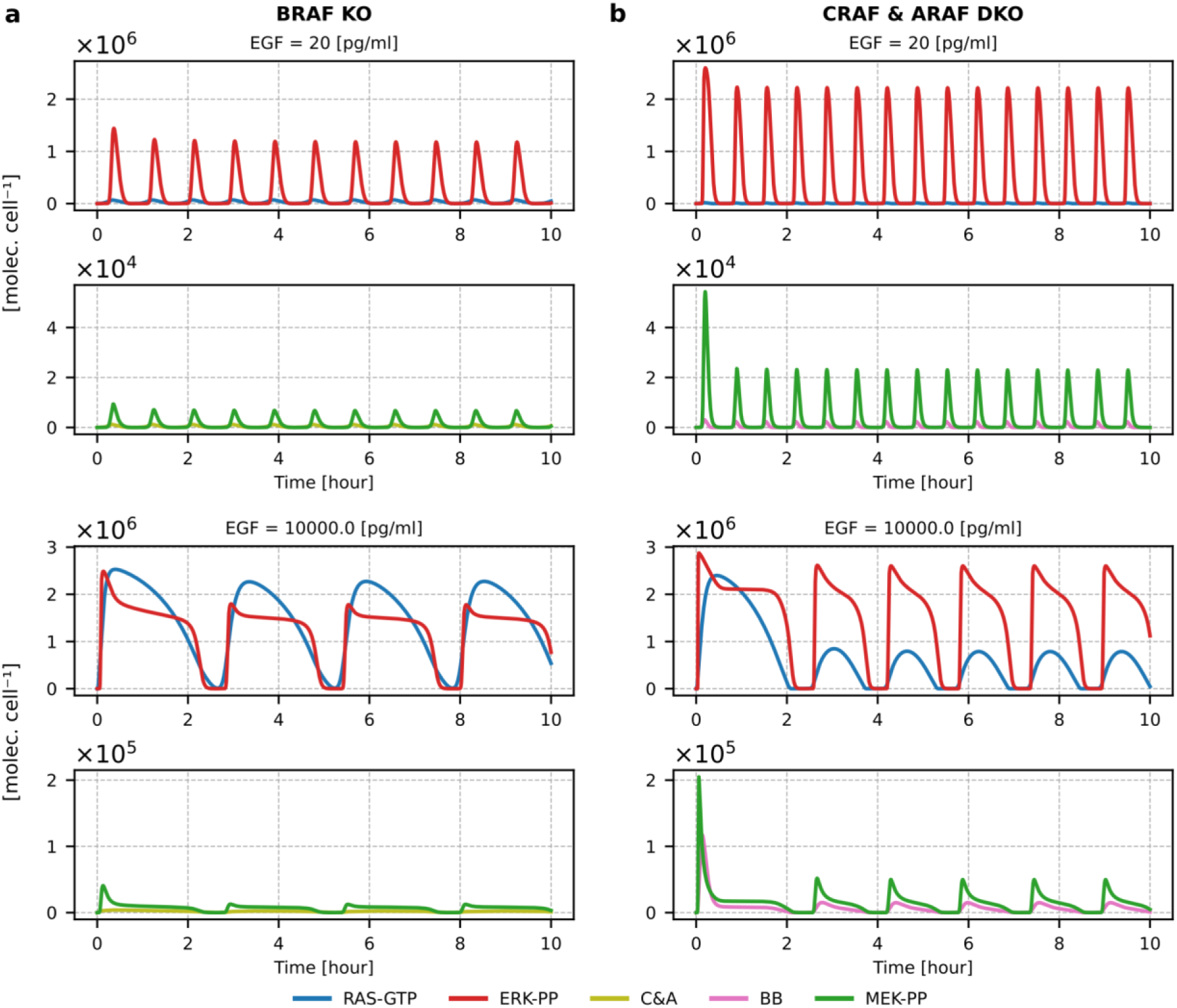
Relaxation oscillations in BRAF KO and ARAF & CRAF DKO cells. **(a)** Profiles of RAS-GTP, C&A dimers, MEKpp, and ERKpp in BRAF KO cells. **(b)** Profiles of RAS-GTP, BB dimers, MEKpp, and ERKpp in ARAF & CRAF DKO cells. Plots are shown for the low and high EGF concentrations; the corresponding plots for the intermediate EGF concentrations, 100 pg/ml and 1000 pg/ml, are provided in Supplementary Figure S2.

The low abundance of C&A dimers may suggest that in BRAF-deficient cells, there will be no ERK activation at all. However, this is not what we observe in the model (Fig. 5a and Supplementary Figure S2a). There are three reasons for this: (1) the positive feedback coupling RAS and SOS is upstream of RAFs and therefore RAS is activated regardless of the presence or absence of BRAF; (2) the number of dimers that do not contain BRAF (i.e., C&A dimers) is much higher in BRAF-deficient cells than in WT cells (compare Fig. 4 and Supplementary Figure S1 with Fig. 5a and Supplementary Figure S2a), because in the latter case, there is no competition between dimers. (3) Finally, the model assumes high rates of MEK and ERK activation, so even a relatively low number of RAF dimers leads to nearly full ERK activation; reducing MEK and ERK activation constants would make the effect of BRAF knockout more pronounced. Nevertheless, experimental data indicate that BRAF inhibitors alone reduce ERK activation only modestly, and combinations of BRAF and MEK inhibitors must be used to inhibit ERK activity^46–48^.

### Switch-like and gradual activation observed at subsequent tiers of MAPK pathway

The presence of a bistable switch that allows for RAS activation on the portion of the membrane and strong signal amplification at the MEK and ERK levels causes ERK activation to have a switch-like character (with the fitted Hill coefficient, *n_H_*, equal 13.4 and EC90/EC10 ratio equal 1.6, while RAS and RAF activation can be characterized as gradual with *n_H_* = 1.1, EC90/EC10 = 84 for RAS and *n_H_* = 2.0, EC90/EC10 = 10.3 for RAF, respectively (Fig. 6a and Table 1). The Hill coefficients have been calculated by fitting the Hill functions to the activation profiles. Of note, the peak and the average level of BRAF-CRAF dimers grow more gradually with EGF concentration than the level of BRAF-BRAF dimers (Fig. 6b and 6d), which makes the interaction of CRAF with ROKα and MST2 change gradually with EGF concentration (Fig. 6c). The level of CRAF competent to bind ROKα increases nearly linearly with *n_H_* = 1.2 and in a broad range (EC90/EC10 = 62); similarly, increases the level of CRAF that is incompetent to bind MST2 (Table 1). These model features are in line with experimental data showing a gradual CRAF activation and switch-like activation of ERK observed in response to serum (FCS) stimulation^14^. Also abundance of CRAF and ROKα complexes grows gradually with the stimulation level^15,43^.

**Figure 6.**
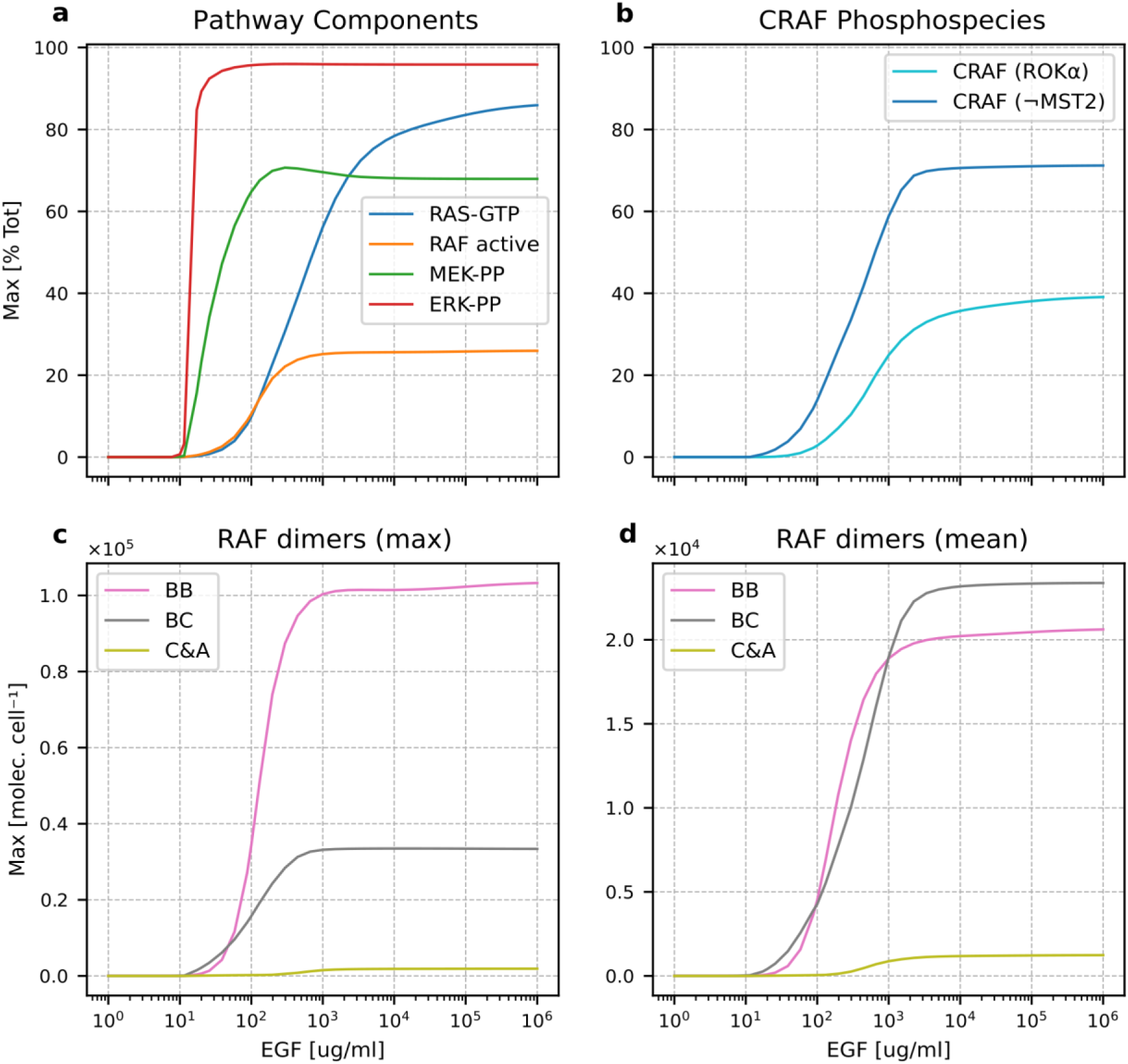
Magnitude of activation of MAPK cascade components as a function of EGF concentration. **(a)** The peak value of RAS-GTP, active RAF, MEKpp, and ERKpp, as a fraction of total protein level. **(b)** The peak value of CRAF (as a fraction of total protein level) in the state in which it can form complexes with ROKα and in the state in which it may not form complexes with MST2. **(c)** The peak value of BB, BC, C&A dimers. **(d)** The mean value over the first 10 hours of EGF stimulation of BB, BC, and C&A dimers.

**Table 1.** Parameters characterizing dose response curves (peak value with respect to EGF concentration) at five tiers of the MAPK pathway. Hill coefficients and EC10, EC50, EC90 values in pg/ml were obtained by fitting the Hill Function.

| Observable | Hill coeff. | EC10 | EC50 | EC90 | EC90/EC10 |
| --- | --- | --- | --- | --- | --- |
| EGFR–SOS | 2.4 | 38 | 109 | 265 | 7.0 |
| RAS–GTP | 1.1 | 90 | 533 | 7564 | 84 |
| RAF active | 2.0 | 38 | 118 | 397 | 10.3 |
| MEKpp | 2.7 | 13.8 | 25 | 78 | 5.7 |
| ERKpp | 13.4 | 11.9 | 14.6 | 18.9 | 1.6 |
| CRAF ROKα-competent | 1.2 | 121 | 634 | 7460 | 62 |
| CRAF MST2-incompetent | 1.3 | 59 | 328 | 1406 | 24 |
| BB dimers | 2.4 | 54 | 130 | 396 | 7.3 |
| BC dimers | 1.6 | 25 | 106 | 375 | 15.1 |

### RAS-dependent prozone effect

RAS and SOS are coupled by a positive feedback loop that introduces bistable switching to the pathway. Not surprisingly, a change in the total RAS level qualitatively influences the activation of RAF isoforms and, thus, downstream components of the MAPK pathway. Reduction of the total RAS level below some threshold (of about 10^6^ molecules per cell) removes bistability and precludes noticeable RAFs activation (Fig. 7a, 7b). An increase of the RAS level above nominal values causes the pathway to lose oscillatory behavior because the activation of ERK is not sufficient to break the SOS-RAS positive feedback loop and terminate RAS activation. For a very high RAS level (beyond physiological limits), RAF dimerization and activation are also reduced because a very high level of RAS dimers reduces the probability that two RAF monomers bind the same RAS dimer, which is needed for RAF dimerization (Fig. 7b).

**Figure 7.**
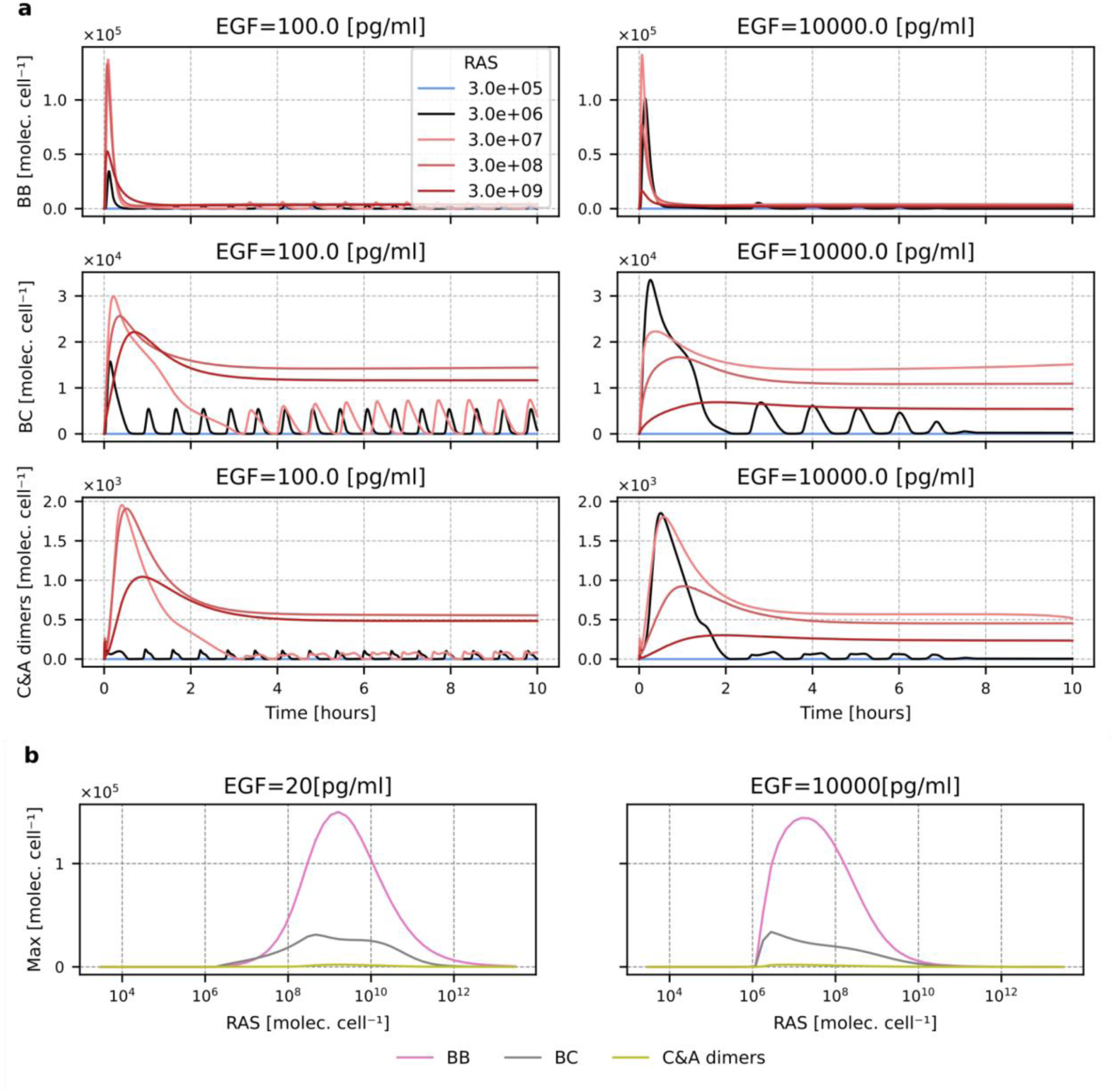
Abundance RAF isoforms dimers as a function of RAS level. **(a)** BB, BC, and C&A dimers time profiles for two EGF concentrations, 100 and 10000 pg/ml, and five RAS levels. The black line corresponds to the nominal RAS level of 3×10^6^ molecules per cell. **(b)** Peak values of BB, BC, and C&A dimers as a function of total RAS for two EGF concentrations, 20 and 10000 pg/ml.

### Dual role of 14-3-3 in RAF isoforms activation

The 14-3-3 dimers play opposing roles in RAF activation. On one hand, they stabilize the closed, autoinhibitory conformation of RAF monomers; when the kinase domain binds the autoinhibitory N-terminal part of the protein, 14-3-3 can simultaneously bind its N- and C-terminal phosphosite, which stabilizes the interaction. Furthermore, 14-3-3 binding to the N-terminal phosphosite interferes with stable binding to RAS, precluding recruitment, dimerization, and activation. On the other hand, 14-3-3 stabilizes RAF dimers by crosslinking RAFs’ C-terminal domains, which promotes RAFs activity. As discussed before, this stabilization effect of 14-3-3 takes place only for dimers that contain BRAF. The picture becomes even more complicated as all three RAF isoforms compete both for binding RAS and for the other RAF isoforms.

First, let us notice that 14-3-3 is required for the system oscillations. When 14-3-3 is absent, or its level is 3×10^5^ (i.e. 10 times lower than the assumed as nominal), we observe only a single peak of active RAF dimers (Fig. 8a). In turn, for EGF concentration equal to 10^4^pg/ml, the higher levels of 14-3-3 (than the nominal) stabilizes oscillations, which for the nominal level of 14-3-3 are damped for this EGF concentration.

**Figure 8.**
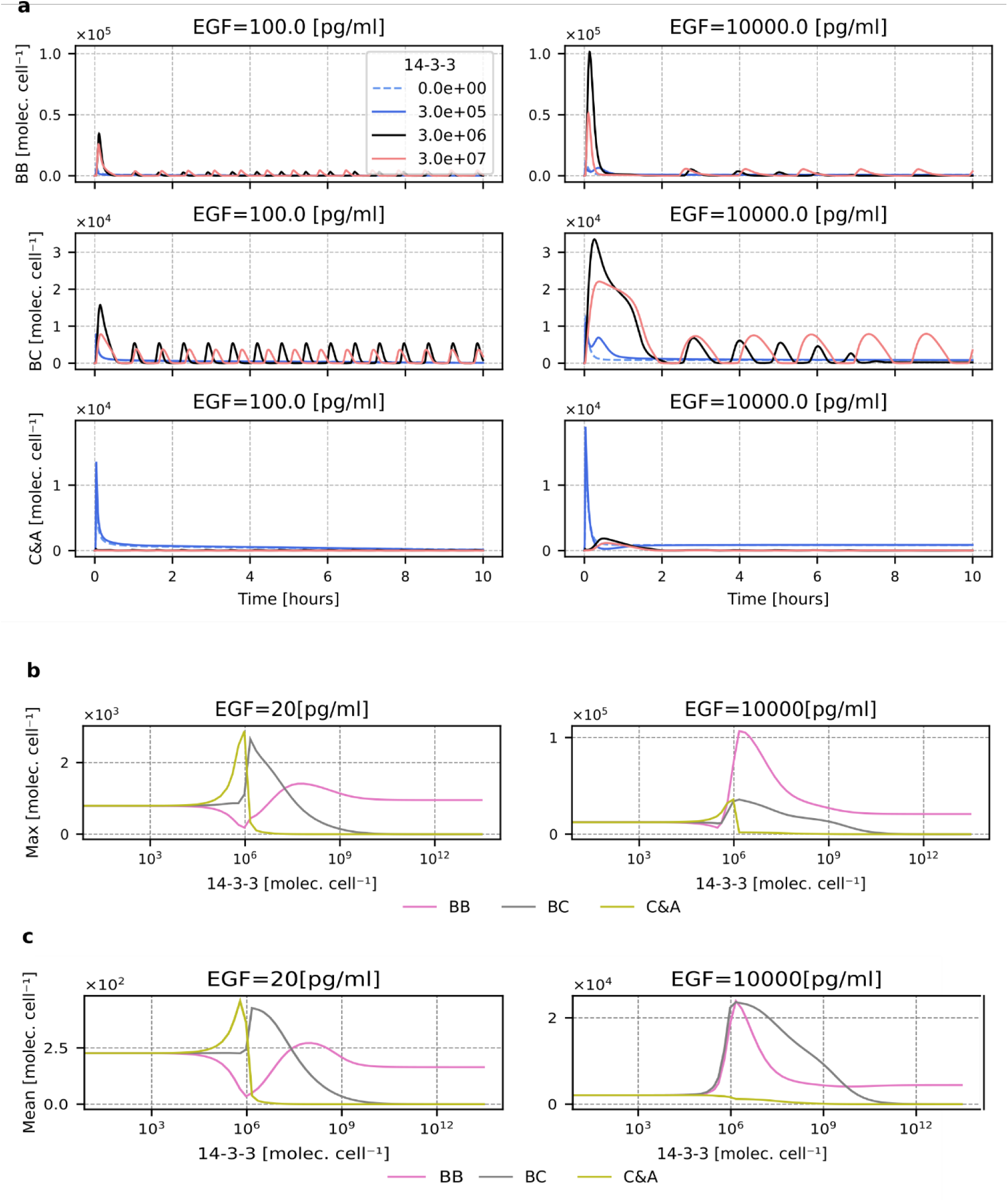
Abundance RAF isoforms dimers as a function of 14-3-3 level. **(a)** BB, BC, and C&A RAF dimers time profiles for two EGF concentrations, 100 and 10000 pg/ml, and four 14-3-3 levels. The black line corresponds to the nominal 14-3-3 level of 3×10^6^ molecules per cell. **(b)** The peak value of BB, BC, and C&A dimers as a function of 14-3-3 for the low and high EGF concentration, 20, 10000 pg/ml. **(c)** The average value of BB, BC, and C&A dimers (in the first 10 hours of EGF stimulation) as a function of 14-3-3 for the low and high EGF concentration, 20, 10000 pg/ml.

As one could expect, the opposing effects of 14-3-3 on RAF activation, and 14-3-3 level-dependent competition between RAF monomers, lead to complex and highly nonlinear dependence of levels of RAF dimers on 14-3-3 abundance. In Fig. 8b and 8c and Supplementary Figure S3A and S3B we show the dependence of peak levels and average level of RAF dimers to 14-3-3 level. First, let us notice that when the level of 14-3-3 is small (so their effect on RAFs is negligible), regardless of the EGF concentration, the peak value of BRAF-BRAF, BRAF-CRAF, and C&A RAF dimers are equal. This is because the only differences between ARAF, BRAF, and CRAF isoforms within our model are associated with their interactions with 14-3-3, and we also assumed that ARAF and CRAF are equally abundant, while BRAF is twice as abundant.

An increase of 14-3-3 is associated with increases in C&A dimers, which is surprising as these dimers are not stabilized by 14-3-3. As shown in Fig. 9a and Supplementary Figure S4a, this effect is caused by BRAF. In BRAF-deficient cells, regardless of EGF concentration, the peak level of C&A dimers decreases monotonically with the number of 14-3-3 molecules (which stabilize ARAF and CRAF monomers in closed conformation). In WT-type cells, 14-3- 3 inhibits BRAF in close conformation, which favors ARAF and CRAF in competition for RAS, which promotes the formation of C&A dimers. This explains why the relative growth of C&A dimers’ peak level is higher for low EGF concentrations when the available pool of active RAS is smaller.

**Figure 9.**
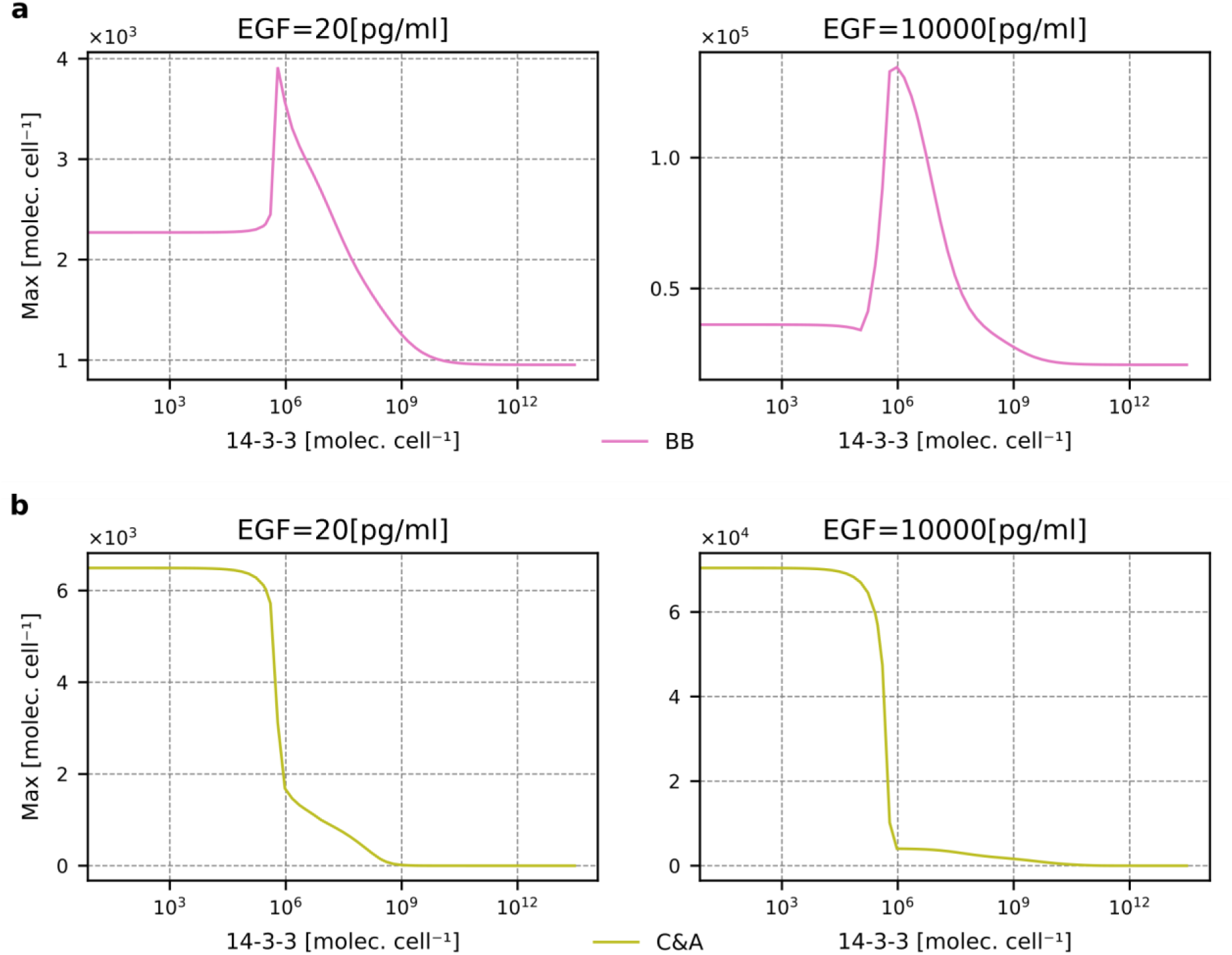
Formation RAF isoforms dimers as a function of 14-3-3 level for BRAF KO and CRAF & ARAF DKO cells. **(a)** The peak value of C&A dimers in BRAF KO cells as a function of 14-3-3 for the low and high EGF concentration 20, 10000 pg/ml. **(b)** The peak value of BB dimers in CRAF & ARAF DKO cells as a function of 14-3-3 for the low and high EGF concentration 20, 10000 pg/ml. The plots corresponding to these in (**a**) and (**b**), but for EGF concentrations of 100, 1000 pg/ml are shown in Supplementary Figure S4.

An increase of C&A dimer level is accompanied by a decrease in BRAF-BRAF dimer level. Again, this decrease is a consequence of competition with two other isoforms – in ARAF and CRAF DKO cells, the decrease of BRAF-BRAF dimers is nearly absent (Fig. 9b and Supplementary Figure S4b). In ARAF and CRAF DKO cells, we may observe that the growth of 14-3-3 level leads first to an increase of BRAF-BRAF level (due to dimers stabilization via 14-3-3), then to a decrease of BRAF-BRAF level due to inhibition of BRAF monomers in the closed form. Let us notice that the strength of dimers stabilization only initially increases with the level of 14-3-3 – for a very high level of 14-3-3, each of the BRAFs forming in a dimer can be loaded with its ‘own’ 14-3-3, which precludes dimer stabilization. The same effect is not observed in the case of monomer stabilization, as 14-3-3 cannot be bound solely to the secondary site.

### Pathway regulation by the MEK isoforms

The ratio of MEK1 to MEK2 determines the strength of the negative feedback from ERK to this tier of the pathway, since MEK1 is the only MEK isoform that possesses an ERK phosphorylation site (Thr292) that downregulates its activity. To the extent that specific functions can be ascribed to individual negative feedback loops, in the context of the current model, the function of the ERK to MEK feedback is to control the shape of the ERK activity pulse. The ablation of this feedback by setting the MEK1/MEK2 (*f*_M1_) ratio equal to zero (Fig. 10), or reducing it by lowering 10-fold the feedback strength (Supplementary Figure S5A) leads to an almost square-like activity profile, representing the maximal possible level of ERK activity. Increasing feedback strength leads to spike-like pulses of activity for low EGF doses and extended pulses with low ERK activity (Supplementary Figure S5A). This latter effect can be understood as follows: reduction of MEK activity immediately reduces ERK activity. This reduces ERK influence on SOS (as well as on RAFs), and thus time to switch off SOS is extended, and consequently, we observe longer ERK activity pulses. These extended pulses (observed for increased strength of ERK to MEK feedback are associated with longer pulses of CRAF in ROKα competent state (Supplementary Figure S5B) and MST2 incompetent state (Supplementary Figure S5C).

**Figure 10.**
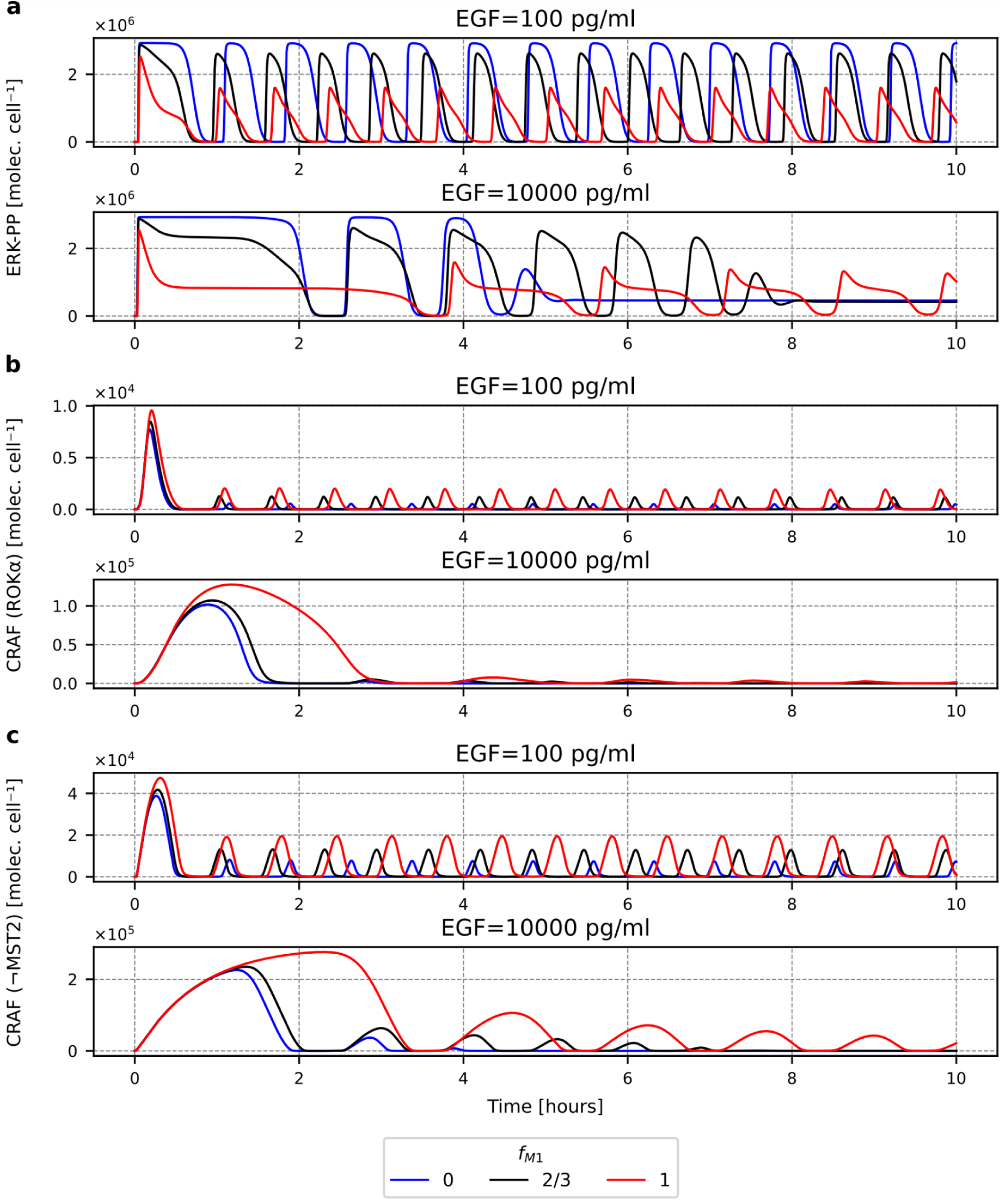
The impact of the MEK1/MEK2 ratio on time profiles of ERKpp, CRAF in ROKα-compatible state, and CRAF in MST2-incompatible state. Black lines correspond to the nominal MEK1/MEK2 ratio *f*_M1_ of ⅔, blue and red lines correspond, respectively, to MEK1 and MEK2 deficiencies compensated by an increase in the abundance of the second isoform. The corresponding plots for different strengths of ERK to MEK1 feedback are provided in Supplementary Figure S5.

## Discussion

We have constructed a rule-based model of the MAPK/ERK pathway to investigate the system’s kinetics. The focus was on regulatory feedback loops, RAF isoforms, and their interaction with 14-3-3 proteins. Instead of fitting to particular cell line responses, we explored the parameter space and RAF and MEK isoform knockouts to investigate the repertoire of responses to growth factor stimulation.

The model accounts for one positive and three negative feedback loops. The positive feedback loop involves SOS and RAS and is responsible for the system’s bistability observed upon silencing the negative feedback loops. All three negative feedback loops emanate from ERK, but have their specific roles.

1. The feedback loop involving SOS is dynamically most important. It encompasses the positive feedback loop, which allows it to frustrate the system’s bistability and induce relaxation oscillations. Consequently, pulsatile and constant-level growth factor stimulation are converted into pulsatile ERK responses.
2. Feedback ERK phosphorylation of RAF disrupts RAF dimers, terminating MEK signaling but enabling further interaction of CRAF monomers with ROKα, promoting cell motility, and with ASK1 inhibiting apoptosis
3. The proximal ERK-MEK feedback functions to shape ERK activity pulses. Its strength increases with the MEK1/MEK2 ratio.

To reconcile nearly all-or-nothing ERK activation with a gradual activation of RAFs, we proposed that at low and intermediate EGF concentrations, growth factor receptors are activated on a portion of the cellular membrane. Consequently, the positive feedback between SOS and RAS induces switching of RAS-GDP to RAS-GTP locally on the cell membrane. Activation of a small proportion of cellular RAS leads to activation of a small proportion of RAFs. This is, however, sufficient to trigger nearly full ERK activation because of ultrasensitivity associated with double MEK and ERK phosphorylation. The proposed mechanism allows MAPK pathways to mediate simultaneously graded responses at the RAF level and switch-like responses at the ERK level. Ultrasensitive, all-or-nothing ERK activation enables temporal pathway inactivation in response to both weak and strong signals. We demonstrated that the different activation patterns of RAF isoforms, and thus RAF dimers, can be attributed to their distinct regulation by 14-3-3. The dimers of 14-3-3 stabilize both the closed, inactive conformation of RAF monomers and the catalytically active conformation of RAF dimers containing BRAF. The specificity of BRAF is caused by the fact that its primary 14-3-3 binding site is on its C-terminal, and therefore (in contrast to CRAF and ARAF) it may bind RAS-GTP with the N-terminal site, without dissociating from 14-3-3. Consequently, it may donate 14-3-3 to stabilize BC and BA dimers. BB homodimers can also be stabilized after 14- 3-3 dissociates from one of the monomers, allowing the remaining one to crosslink the dimer. CRAF and ARAF may bind RAS-GTP only after 14-3-3 dissociates, and thus, RAF dimers that do not contain BRAF are not stabilized by 14-3-3, and their abundance is much lower. Nevertheless, the model predicts that although in WT cells, BRAF containing dimers dominate, the effect of BRAF knockout on ERK activity is weak. This is because (1) SOS-RAS positive switch is independent of RAF signaling, (2) BRAF deficiency leads to an increase of C&A dimers, and (3) ultrasensitivity at MEK and ERK level allows for nearly full ERK activation at low levels of active RAFs.

The fact that 14-3-3 stabilizes both inactive RAF monomers and active RAF dimers (containing BRAF) causes the abundance of specific RAF dimers to be a non-monotonic function of the 14-3-3 level. The complexity of the picture is further increased by the competition between RAF isoforms for RAS-GTP (important in the case of low EGF concentrations). Consequently, the model predicts qualitatively different dependence of BC, BB, and C&A dimers on the level of 14-3-3. For the nominal level of 14-3-3, the peak value of BB dimers is higher than that of BC and C&A dimers for most, but very low EGF concentrations, for which BC dimers dominate. When the average level over the first 10 hours of EGF stimulation is considered, BC dimers dominate the pool of RAF dimers.

The relative abundance of RAF dimers and engaged RAF monomers is not important for global RAF kinase activity, but becomes crucial when the interaction of CRAF and ARAF with other proteins is considered. Engagement of ARAF and CRAF in RAF dimers releases MST2, allowing it to participate in the proapoptotic Hippo pathway (Rauch ARAF). The model indicates that ARAF and CRAF fraction, capable of binding MST2, decreases gradually with EGF concentration, meaning that strong, long-lasting EGF signals can be proapoptotic^14^. CRAF is released from BC dimers in the open conformation in which it can bind ROKα, promoting cell motility, and coordinating it with proliferation^15,43^.

In summary, the constructed model indicates that the presence of RAF and MEK isoforms, together with the regulatory features of the MAPK/ERK pathway, allows for a broad repertoire of dynamic responses to growth factor stimulation. This implies that different cell types, having specific proportions of signaling proteins, may exhibit qualitatively different responses, employing the MAPK/ERK pathway to divers physiological functions.

## Methods

### Rule-based modeling

The model was implemented in a rule-based paradigm, using BioNetGen language (BNGL) within the BioNetGen environment (the code provided as Supplementary Code S1)^37,38,49^. BNGL models are formulated by enumerating rules, which encode possible reactions. A set of rules specifying a model is then automatically processed to generate the corresponding system of coupled ODEs; the system is assumed to follow mass-action kinetics. The systems of ODEs were numerically integrated using solvers provided with BNGL (CVODE) at default settings.

### Model description

The model follows the established basic growth factor signal transduction scheme in the RAF/ERK cascade. Briefly, EGF binds and activates the EGFR receptor, which can then bind SOS. EGFR-bound SOS subsequently activates RAS. Activated RAS binds and activates RAF isoforms. Activated RAF subsequently phosphorylates and activates MEK1/2, which does the same to ERK1/2. ERK1/2 phosphorylates SOS, RAF isoforms, and MEK1, disrupting their activity and forming negative feedback loops. The positive feedback between SOS and RAS arises since RAS-GTP allosterically binds to SOS, promoting the conversion of RAS-GDP to RAS-GTP.

The model was implemented in BNGL due to its significant inherent combinatorial complexity. It comprises 11 proteins: EGFR, SOS, RasGAP, RAS, BRAF/CRAF/ARAF isoforms, 14-3-3, MEK1 and MEK2 isoforms, and ERK. It has been specified in BNGL setting 122 reaction rules and parameters (provided in Supplementary Text S1), generating a network of 9088 chemical reactions and 609 molecular species represented by the corresponding system of 609 coupled ODEs.

#### The RAF Isoforms

The model accounts for all three known vertebrate RAF isoforms: ARAF/BRAF/CRAF (RAF1). Their structure and regulation have been implemented in significant mechanistic detail. Each isoform follows the structure of a generic RAF protein, featuring the N- and C-terminal 14-3-3 binding sites, the N-terminal acidic domain (NtA), and the dimerization interface DIF. The RAF isoforms are known to possess the inactive “closed” and active “open” conformations. In the “closed” conformation, the C-terminal catalytic domain-containing segment of the protein is occluded by the N-terminal autoinhibitory segment; this conformation is further stabilized by the intramolecular crosslinking by a 14-3-3 protein bound at the N-terminal and C-terminal phosphosites. In contrast, in the active “open” conformation, the interaction between the N- and C-terminal segments is disrupted, allowing for the interaction between the catalytic domain and substrates. This conformation is likely stabilized by the phosphorylation of the NtA domain, and this phosphorylation is used as the proxy for “open” active RAF. The RAF isoforms share the same activation scenario:

1. Dimerized RAS-GTP recruits RAF protein monomers. For this interaction to occur, the N-terminal 14-3-3 binding site must be unoccupied
2. The two RAF proteins bound to the RAS-GTP dimer subsequently dimerize. The RAF dimers containing BRAF are stabilized by the intramolecular crosslinking via the 14-3- 3 bound to the C-terminal
3. The dimerization triggers the autophosphorylation of the NtA domains of the constituent RAF protomers. Such a dimer is considered to be catalytically active.
4. The RAF dimer detaches itself from the RAS-GTP dimer.
5. ERK phosphorylates the DIF of the RAF protein, triggering the breakdown of the dimers and terminating signaling.

While the RAF isoforms share the same overall structure, there are discernible differences between them in terms of activity and regulation, hinting at specialized functions. The significant difference between them is that in BRAF, the high-affinity 14-3-3 binding site is the C-terminal one, unlike the N-terminal one in ARAF and CRAF. As a consequence, the ARAF and CRAF proteins bound to RAS-GTP are devoid of 14-3-3, while BRAF can maintain its interaction with 14-3-3 via its C-terminal site. This favors stability of BRAF heterodimers with other isoforms, as BRAF can act as a donor of 14-3-3 to the C-terminal binding site of the other isoform within the dimer, which is likely to be unoccupied and thus lends itself to crosslinking and dimer stabilization. In the case of BRAF homodimers, however, the simultaneous occupancy of both C-terminal binding sites by two distinct 14-3-3 proteins can interfere with crosslinking and stabilization. ARAF and CRAF homodimers and CRAF-ARAF heterodimers are not crosslinked by 14-3-3, which effectively reduces their stability. This explains the relative abundance of BRAF-containing heterodimers that accounts for the observation that the BRAF heterodimers (BRAF-CRAF in particular) are the main activators of MEK1/2.

#### Nested Feedback Architecture

It has been experimentally observed that the RAF/ERK cascade is capable of producing ppERK pulses with a nearly EGF concentration-independent amplitude but an EGF concentration-dependent period. This suggests frequency-based rather than typically assumed amplitude-based encoding of the signal strength. We previously demonstrated that the RAF/ERK cascade could exhibit this behavior if it were capable of relaxation oscillations^32^. Relaxation oscillations involve periodic and abrupt alternations between two stable equilibrium states, resulting in non-sinusoidal repetitive patterns such as spikes. This behavior emerges from the activity of two opposing but coupled feedback processes with significantly differing time scales, whereupon the slow-acting negative feedback loop encloses the fast-acting positive feedback loop. We previously identified such a motif in the structure of the RAF/ERK cascade, demonstrated that it can produce the observed frequency-based encoding, and now incorporate it into the current model.

The outer slow-acting negative loop is based on the negative feedback phosphorylation of SOS by ppERK. SOS contains four sites targeted by ERK, and phosphorylation of any of them precludes its binding to activated EGFR receptors and membrane recruitment. This forms the longest and correspondingly slowest feedback loop in the RAF/ERK cascade. SOS is also the target of the inner fast-acting positive feedback loop. SOS undergoes positive regulation by its activated substrate RAS-GTP. In particular, RAS-GTP can bind SOS through a separate domain (REM) and allosterically increase its GEF activity, resulting in positive feedback. This feedback loop is fast due to its short range. In summary, SOS is a nexus of a long, slow-acting negative feedback loop and the enclosed short, fast-acting positive feedback loop, which produces relaxation oscillations.

#### Signaling Scenario

In unstimulated cells, RAS is in the GDP-bound monomeric state on the plasma membrane, while SOS is in the cytoplasm in its inactive form and unphosphorylated on its ERK feedback sites. The ARAF, BRAF and CRAF isoforms molecules are in the inactive “closed” conformation and are phosphorylated on their N-terminal (ARAF – Ser214, BRAF-Ser365, CRAF – Ser259) and C-terminal (ARAF – Ser582, BRAF-Ser729, CRAF – Ser621) 14-3-3 binding sites, which enables further “locking” of the conformation by 14-3-3 crosslinking. They are also unphosphorylated on their respective NtA domains (ARAF – Ser299, CRAF – Ser338). In contrast, BRAF is assumed to be constitutively phosphorylated on its NtA site (Ser446). It is assumed that, in contrast to ARAF and CRAF, BRAF may be crosslinked by 14- 3-3 when phosphorylated at Ser446. All RAF isoforms are unphosphorylated on their DIF region. MEK and ERK are inactive, unphosphorylated on their activation loops and feedback sites.

Upon stimulation with the ligand, EGFR becomes activated and recruits SOS to the membrane. The rate of EGFR activation corresponds to the concentration of EGF scaled by the fraction of the membrane *f* undergoing activation, *f* = *EGF*/(*EGF +* M_EGF_), with MEGF = 300 pg/ml, i.e., we assume that at low EGF concentration only part of the membrane becomes active. SOS subsequently recruits RAS-GDP via its GEF domain and induces RAS to exchange GDP for GTP, which leads to its activation and dimerization. SOS can also independently bind RAS-GTP via the REM domain, which allosterically upregulates its GEF activity. A RAS-GTP monomer can be recruited and deactivated by RasGAP by inducing transition from RAS-GTP back RAS-GDP.

Dimerized RAS-GTP recruits two RAF monomers; to be competent for RAS recruitment RAF isoform molecules have to be free of 14-3-3 at their N-terminal binding sites, which enables interaction with RAS. The bound RAF proteins can subsequently dimerize on the RAS-GTP dimer platform via their DIFs. The dimers containing BRAF may be stabilized by 14-3-3 crosslinking via the C-terminal 14-3-3 binding sites. RAF dimers subsequently detach from the RAS dimers. RAF dimers are considered catalytically active and phosphorylate MEK1/2 on their activation loops. MEK1/2 consequently activate ERK by phosphorylating its activation loop.

Phosphorylated ERK can disrupt the activity of upstream components, forming negative feedback loops. We have accounted for the following:

1. ERK phosphorylates four sites on SOS, each individually sufficient to prevent the interaction between EGFR and SOS. This loop is a part of the relaxation oscillation motif.
2. ERK phosphorylates RAF monomers and protomers on the DIF region, blocking dimerization and disrupting existing dimers.
3. ERK phosphorylates MEK1 at Thr292, which (by recruiting an implicit phosphatase/phosphatase-containing complex) facilitates dephosphorylation of the MEK activation loop.

The negative feedback to the RAF dimers plays a relevant role, independent of pathway deactivation. Specifically, ARAF and CRAF monomers produced from disrupted dimers retain their phosphorylation of their NtA regions; this keeps them in the “open” conformation and enables interaction with ROKα.

## Acknowledgements

This work was supported by the National Science Centre, Poland (https://ncn.gov.pl) grant 2019/35/B/NZ2/03898 to T.L. The funder had no role in study design, data collection and analysis, decision to publish, or preparation of the manuscript.

## Author Contributions

Conceived the study: P.K. and T.L. Formulated and analyzed mathematical models: P.K. Interpreted the results: P.K. and T.L. Wrote the manuscript: P.K. and T.L.

## Data Availability Statement

Data supporting this study are included within the article and/or supporting materials.

## Supplementary Material includes

Supplementary Text S1 (appended to main text),

Supplementary Figures S1–S5 (appended to main text),

Supplementary Code S1 (zip file).

## Supplementary Text S1

### Species definitions

**EGFR(egf∼I∼A, sos)** – EGFR, EGF receptor

● **egf:** indicates the inactive (I) or active (A) state of the receptor
● **sos:** the SOS binding domain, active receptor can bind SOS

**SOS(egfr,rem,S∼P0∼P1∼P2∼P3∼P4) –** SOS, a guanine nucleotide exchange factor (GEF), activates RAS-GDP by inducing nucleotide exchange from GDP to GTP and producing RAS- GTP

● **egfr**: the EGFR-binding site
● **rem**: a secondary RAS biding domain, when occupied by RAS it allosterically enhances the GEF activity of SOS
● **S**: a flag to indicate the number of phosphorylated inactivating sites, SOS possesses four such sites, which are subject to distributive and independent feedback phosphorylation by ERK1/2; phosphorylation of any of them precludes SOS-EGFR binding.

**RasGAP(ras) –** RAS GTP-ase Activating Protein, which inactivates RAS-GTP by inducing the hydrolysis of GTP to GDP:

● **ras:** the RAS binding domain

**RAS(sos,dim,raf,nt∼GDP∼GTP) –** RAS, a small GTPase, binds and recruits RAF isoforms to the plasma membrane and initiates their activation:

● **sos**: the SOS binding site
● **dim**: the dimerization interface
● **raf**: the RAF binding site
● **nt**: a flag to indicate the activation status of RAS. RAS can be loaded either with GDP (**nt∼GDP** – considered inactive) or GTP (**nt∼GTP** – considered active). GDP and GTP are not explicitly represented.

**RAF(iso∼B∼C∼A,ras,N_ftt∼P0∼P1,dif∼P0∼P1,NtA∼P0∼P1,C_ftt∼P0∼P1) –** a RAF kinase with BRAF,CRAF, and ARAF isoforms

Due to the regulatory and structural similarities between RAF isoforms we introduced a generalized RAF isoform species. When an isoform deviates in its regulation from the assumed baseline, it is explicitly identified in the corresponding rules.

● **iso:** specifies the identity of the isoform (B/C/A) if required
● **ras:** the RAS binding site, collapses both the RBD and CRD domains
● **N_ftt:** the N-terminal 14-3-3 binding site; ARAF – Ser^214^, CRAF – Ser^259^ BRAF – Ser^365^
● **dif:** the dimerization interface/domain
● **NtA:** the N-terminal acidic domain whose phosphorylation promotes the active conformation and protein-protein interactions; ARAF – Ser^299^, CRAF – Ser^338^, BRAF – Ser^446^
● **C_ftt:** the C-terminal 14-3-3 binding site, ARAF – Ser^582^,CRAF – Ser^621^, BRAF – Ser^729^

**MEK(iso∼mek1∼mek2,dim,NFS∼P0∼P1,al∼P0∼P1∼P2) –** MEK kinase with MEK1 and MEK2 isoforms, ERK-activating kinases

● **iso:** the identity of the isoform (MEK1 or MEK2)
● **dim:** the dimerization interface
● **NFS:** the ERK negative feedback site. When MEK is phosphorylated at this site it is rapidly dephosphorylated on its activation loop (**al**). Within a dimer, the phosphorylation of one protomer at this site confers the rapid dephosphorylation of the activation loop of the second protomer. This site is functional only in MEK1 (T292)
● **al:** the activation loop, biphosphorylated (**al∼P2**) MEK1/2 is catalytically activated

**ERK(al∼P0∼P1∼P2) –** ERK, the last (output) kinase in the canonical ERK cascade, mediates negative feedback to SOS, A/B/CRAF, and MEK

● **al**: the activation loop, when biphosphorylated (**al∼P2**) ERK is catalytically activated

**FTT(raf,dim,iso∼A∼B) –** 14-3-3 protein, binds A/B/CRAF

● **raf:** a RAF isoform binding site. 14-3-3 can bind ARAF on pSer^214^ and pSer^582^, BRAF on pSer^356^ and pSer^729^, and CRAF on pSer^259^ and pSer^621^
● **dim:** the dimerization interface with other 14-3-3 monomers
● **iso:** specifies the minimal set of two distinct isoforms to account for homo- and heterodimeric nature of 14-3-3 proteins

### RAF isoforms binding sites

| Isoform | N-terminal 14-3-3 binding site | C-terminal 14-3-3 binding site | NtA domain | DIF negative feedback sites |
| --- | --- | --- | --- | --- |
| ARAF | Ser214 | Ser582 | Ser299 | Not mapped |
| BRAF | Ser365 | Ser729 | Ser446 | Thr401, Ser750, Tyr753 |
| CRAF | Ser259 | Ser621 | Ser338 | Ser289/296/301 |

### Parameters

#### 1. Molecular Species

Parameters 1. Molecular Species
| Concentrations |  |  |
| --- | --- | --- |
| Parameter | Value | Description |
| *EGFR_tot* | $3 \times 10^5 \text{ [molec} \times \text{cell}^{-1}]$ | EGFR receptor |
| *SOS_tot* | $1 \times 10^4 \text{ [molec} \times \text{cell}^{-1}]$ | SOS (all isoforms) |
| *RasGAP_tot* | $1 \times 10^5 \text{ [molec} \times \text{cell}^{-1}]$ | RAS-GAP |
| *RAS_tot* | $3 \times 10^6 \text{ [molec} \times \text{cell}^{-1}]$ | RAS (all isoforms) |
| *BRAF_tot* | $6 \times 10^5 \text{ [molec} \times \text{cell}^{-1}]$ | BRAF |
| *CRAF_tot* | $3 \times 10^5 \text{ [molec} \times \text{cell}^{-1}]$ | CRAF |
| *ARAF_tot* | $3 \times 10^5 \text{ [molec} \times \text{cell}^{-1}]$ | ARAF |
| *MEK_tot* | $3 \times 10^5 \text{ [molec} \times \text{cell}^{-1}]$ | MEK1 and MEK2 |
| *ERK_tot* | $3 \times 10^6 \text{ [molec} \times \text{cell}^{-1}]$ | ERK (both isoforms) |
| *FTT_tot* | $3 \times 10^6 \text{ [molec} \times \text{cell}^{-1}]$ | 14-3-3 |

Wpisz tutaj równanie.

#### 2. Kinetic Parameters

| Quiescent state and signal-independent interactions |  |  |
| --- | --- | --- |
| Parameter | Value | Description |
| *p_S365* | $1 \times 10^{-1} \text{ [s}^{-1}]$ | Phosphorylation of BRAF-S365 |
| *q_S365* | $1 \times 10^{-1} \text{ [s}^{-1}]$ | Dephosphorylation of BRAF-S365p |
| *b_BF_p729* | $3 \times 10^{-8} \text{ [molec} \times \text{s}^{-1}]$ | Binding of 14-3-3 to BRAF-S729p |
| *u_BF_p729* | $3 \times 10^{-2} \text{ [s}^{-1}]$ | Dissociation of 14-3-3 from BRAF-S729p |
| *p_NS_AC* | $3 \times 10^{-1} \text{ [s}^{-1}]$ | Phosphorylation of CRAF-S259 and ARAF-S214 |
| *q_NSp_AC* | $1 \times 10^{-2} \text{ [s}^{-1}]$ | Dephosphorylation of CRAF-S259p and ARAF-S214p |
| *p_CS_AC* | $1 \times 10^{-2} \text{ [s}^{-1}]$ | (Auto)Phosphorylation of CRAF-S621 and ARAF-S582 |
| *p_CS_AC_dim* | $100 \text{ [s}^{-1}]$ | Dimer induced (auto)phosphorylation of CRAF-S621 and ARAF-S582 |
| *q_CS_AC* | $3 \times 10^{-1} \text{ [s}^{-1}]$ | Dephosphorylation of CRAF-S621p ARAF-S582p |
| *b_FNp_AC* | $1 \times 10^{-6} \text{ [molec} \times \text{s}]^{-1}$ | Binding of 14-3-3 to CRAF-S259p and ARAF-S214p |
| *u_FNp_AC* | $1 \text{ [s}^{-1}]$ | Dissociation of 14-3-3 from CRAF-S259p and ARAF-S214p |
| *b_RF_intra* | $100 \text{ [s}^{-1}]$ | Intramolecular crosslinking of A/B/CRAF by 14-3-3 |
| $u_{RF\_inter}$ | $3 \times 10^{-2} [molec \times s]^{-1}$ | Intramolecular decrosslinking of A/B/CRAF by 14-3-3 |
| $b_{RF\_inter}$ | $100 [s^{-1}]$ | Intermolecular crosslinking of RAF dimers by 14-3-3 |
| $u_{RF\_intra}$ | $3 \times 10^{-2} [molec \times s]^{-1}$ | Intermolecular decrosslinking of RAF dimers by 14-3-3 |
| $b_{MEK11}$ | $1 \times 10^{-5} [molec \times s]^{-1}$ | Homodimerization of MEK1 |
| $u_{MEK11}$ | $1 \times 10^{-3} [s^{-1}]$ | Dissociation of MEK1 homodimers |
| $b_{MEK22}$ | $1 \times 10^{-5} [molec \times s]^{-1}$ | Homodimerization of MEK2 |
| $u_{MEK22}$ | $3 \times 10^{-2} [s^{-1}]$ | Dissociation of MEK2 homodimers |
| $b_{MEK12}$ | $1 \times 10^{-5} [molec \times s]^{-1}$ | Heterodimerization of MEK1 and MEK2 |
| $u_{MEK12}$ | $1 \times 10^{-3} [s^{-1}]$ | Dissociation of MEK1/MEK2 heterodimers |

| Signal Transduction - Membrane |  |  |
| --- | --- | --- |
| Parameter | Value | Description |
| $a_{EGFR}$ | $1 \times 10^{-6} \times EGF [s^{-1}]$ | Activation of EGFR by EGF |
| $i_{EGFR}$ | $1 \times 10^{-2} [s^{-1}]$ | Deactivation of EGFR |
| $b_1$ | $1 \times 10^{-5} [molec \times s]^{-1}$ | Binding of unphosphorylated SOS to activated EGFR |
| $u_{1a}$ | $1 \times 10^{-2} [s^{-1}]$ | Dissociation of unphosphorylated SOS from active EGFR |
| $u_{1b}$ | $100 [s^{-1}]$ | Dissociation of unphosphorylated SOS from inactive EGFR |
| $b_{2a}$ | $1 \times 10^{-6} / frac [molec \times s]^{-1}$ | Binding of RAS-GDP to SOS(rem) |
| $u_{2a}$ | $1 [s^{-1}]$ | Dissociation of RAS-GDP from SOS(rem) |
| $b_{2b}$ | $b_{2a} \times 10 [molec \times s]^{-1}$ | Binding of RAS-GTP to SOS(rem) |
| $u_{2b}$ | $1 [s^{-1}]$ | Dissociation of RAS-GTP from SOS(rem) |
| $k_{2a}$ | $1 \times 10^{-6} / frac [s^{-1}]$ | RAS activation via SOS-mediated nucleotide exchange; SOS allosterically upregulated by REM-bound RAS-GDP |
| $k_{2b}$ | $k_{2a} \times 10 [s^{-1}]$ | RAS activation via SOS-mediated nucleotide exchange; SOS allosterically upregulated by REM-bound RAS-GTP |
| $b_3$ | $1 \times 10^{-5} / frac [molec \times s]^{-1}$ | Biding of RAS-GTP to RasGAP |
| $u_3$ | $1 \times 10^{-2} [s^{-1}]$ | Dissociation of RAS-GDP from RasGAP |
| $k_3$ | $100 [s^{-1}]$ | RAS inactivation via RasGAP induced GTP hydrolysis |
| $b_4$ | $1 \times 10^{-5} / \text{frac} [s^{-1}]$ | Homodimerization of SOS-free RAS-GTP monomers |
| $b_1$ | $10^{-1} [s^{-1}]$ | Dissociation of SOS-free RAS-GTP dimers |

| Signal Transduction – RAF/MEK/ERK Cassette |  |  |
| --- | --- | --- |
| Parameter | Value | Description |
| $b_{GR}$ | $3 \times 10^{-5} [molec \times s]^{-1}$ | Binding of A/B/CRAF to RAS-GTP |
| $u_{GR}$ | $100 [s^{-1}]$ | Dissociation of A/B/CRAF from RAS-GDP |
| $u_{GR\_GTP}$ | $1 \times 10^{-1} [s^{-1}]$ | Dissociation of A/B/CRAF from RAS-GTP |
| $u_{GRR\_dim}$ | $10 [s^{-1}]$ | Dissociation of RAF dimers from RAS-GDP |
| $u_{GRR\_dim\_GTP}$ | $1 [s^{-1}]$ | Dissociation of RAF dimers from RAS-GTP |
| $b_{GRR\_dim}$ | $100 [s^{-1}]$ | Homo- and heterodimerization of RAF isoforms on RAS |
| $u_{GRR\_dim\_fast}$ | $3 [s^{-1}]$ | Dissociation of RAF homo- and heterodimers phosphorylated at DIF domain on RAS platform |
| $u_{RR\_dim}$ | $1 [s^{-1}]$ | Dissociation of RAF homo- and heterodimers not phosphorylated at DIF domain in the cytoplasm |
| $u_{RR\_dim\_fast}$ | $3 \times 10^1 [s^{-1}]$ | Dissociation of RAF homo- and heterodimers phosphorylated at DIF domain in the cytoplasm |
| $p_{NtA\_dim\_AC}$ | $1 \times 10^{-1} [s^{-1}]$ | Phosphorylation of CRAF-S338 and ARAF-S299 within dimer |
| $q_{NtA\_cyt\_AC}$ | $1 \times 10^{-1} [s^{-1}]$ | Dephosphorylation of CRAF-S338 and ARAF-S299 in the cytoplasm |
| $p_{RM}$ | $1 \times 10^{-6} [s^{-1}]$ | Phosphorylation of MEK by BRAF or RAF1 on activation loop |
| $p_{M1E}$ | $1 \times 10^{-6} [s^{-1}]$ | Phosphorylation of ERK by MEK1 on activation loop |
| $p_{M2E}$ | $X \times p_{M1E}$ | Phosphorylation of ERK by MEK2 on activation loop |
| $q_{AL\_MEK}$ | $3 \times 10^{-3} [s^{-1}]$ | Dephosphorylation of MEK on activation loop |
| $q_{AL\_ERK}$ | $1 \times 10^{-2} [s^{-1}]$ | Dephosphorylation of ERK on activation loop |
| $q_{MP}$ | $100 [s^{-1}]$ | Dephosphorylation of MEK activation loop after negative feedback phosphorylation |

| Signal Transduction – Negative feedback regulation |  |  |
| --- | --- | --- |
| Parameter | Value | Description |
| $p_{nfSOS}$ | $1 \times 10^{-8} [molec \times s]^{-1}$ | Feedback phosphorylation of SOS by ERK |
| $q_{nfSOS}$ | $1 \times 10^{-3} [s^{-1}]$ | Dephosphorylation of SOS on the feedback sites |
| $p_{nfRAF}$ | $1 \times 10^{-8} [molec \times s]^{-1}$ | Feedback phosphorylation of ARAF/BRAF/CRAF on DIF by ERK |
| $q_{nfRAF}$ | $1 \times 10^{-4} [s^{-1}]$ | Dephosphorylation of ARAF/BRAF/CRAF on the feedback sites |
| $p_{nfMEK}$ | $1 \times 10^{-9} [molec \times s]^{-1}$ | Feedback phosphorylation of MEK1 by ERK |
| $q_{nfMEK}$ | $3 \times 10^{-4} [s^{-1}]$ | Dephosphorylation of MEK1 on its feedback site |

| Coefficients and Auxilliary Parameters |  |  |
| --- | --- | --- |
| Parameter | Value | Description |
| $EGF_{MM}$ | 300 | MM dose coefficient |
| $EGFs$ | Set | Ligand concentration [pg/ml] |
| $EGF$ | Set | Ligand concentration [pg/ml] |
| $Frac$ | $EGFs / (EGFs + EGF_{MM})$ | The fraction of the membrane undergoing activation |
| $X$ | 5 | The ratio MEK1 and MEK2 catalytic activity |
| $f_{M1}$ | 2/3 | The ratio of MEK1 to MEK2 |
| $PF$ | 10 | Allosteric upregulation of (1) RAS binding to SOS and (2) of RAS GEF-activity by RAS bound to SOS via the Rem domain |
| $f_{nfDIM}$ | 30 | Scaling of dimer dissociation due to phosphorylation of DIF |
| $f_{GRR}$ | $1 \times 10^{-1}$ | Scaling of dimer dissociation for dimers bound to RAS platforms |

---

**Figure S1.**
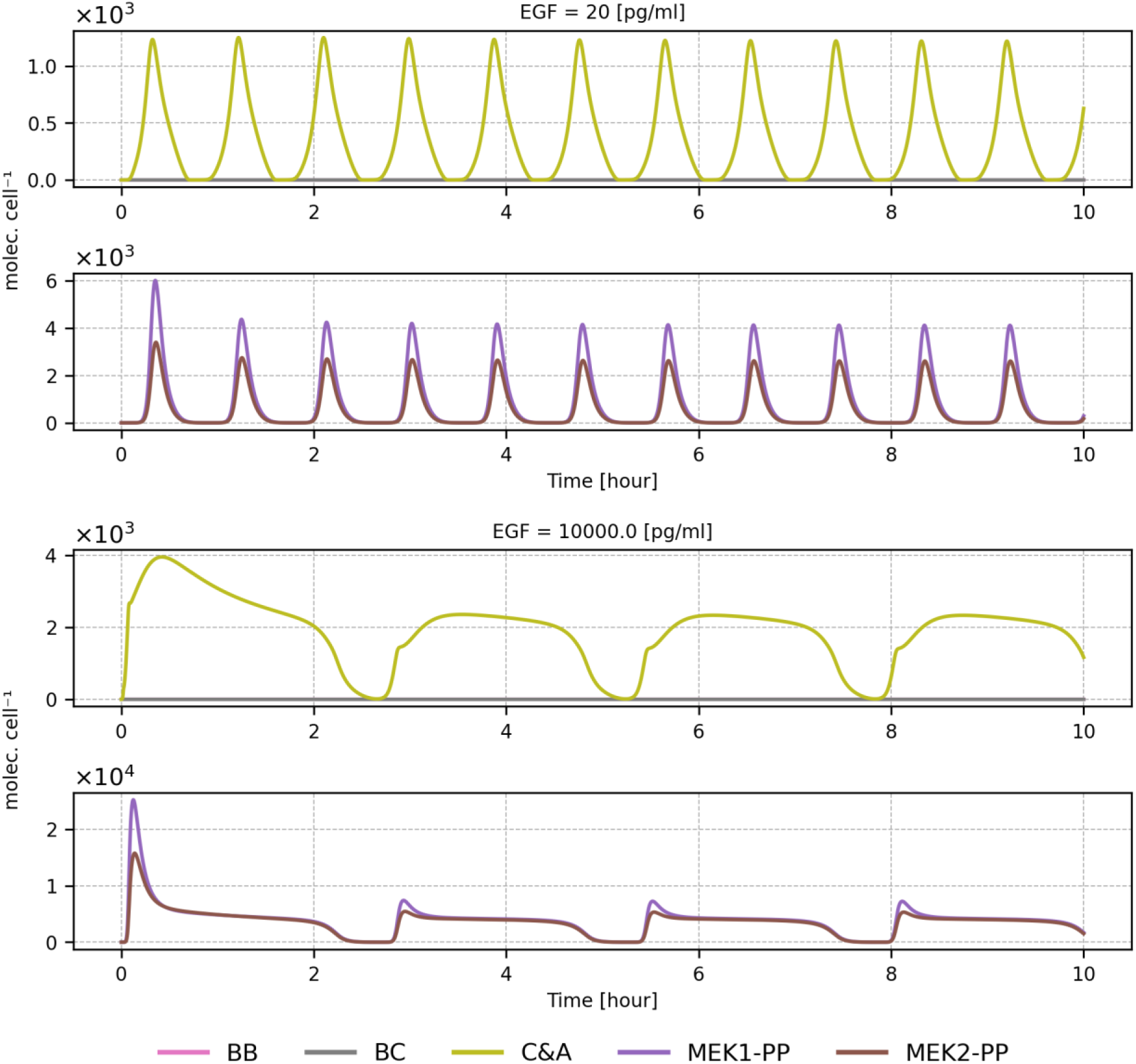
RAF and MEK isoforms time profiles during relaxation oscillations. The plot shows abundances of BB, BC, and C&A RAF dimers, and MEK1pp, MEK2pp molecules for low and high EGF concentrations. C&A dimers stand for all RAF dimers not containing BRAF. The corresponding plots for the low and high EGF concentrations, 20 pg/ml and 10000 pg/ml, are given in Fig. 4.

**Figure S2.**
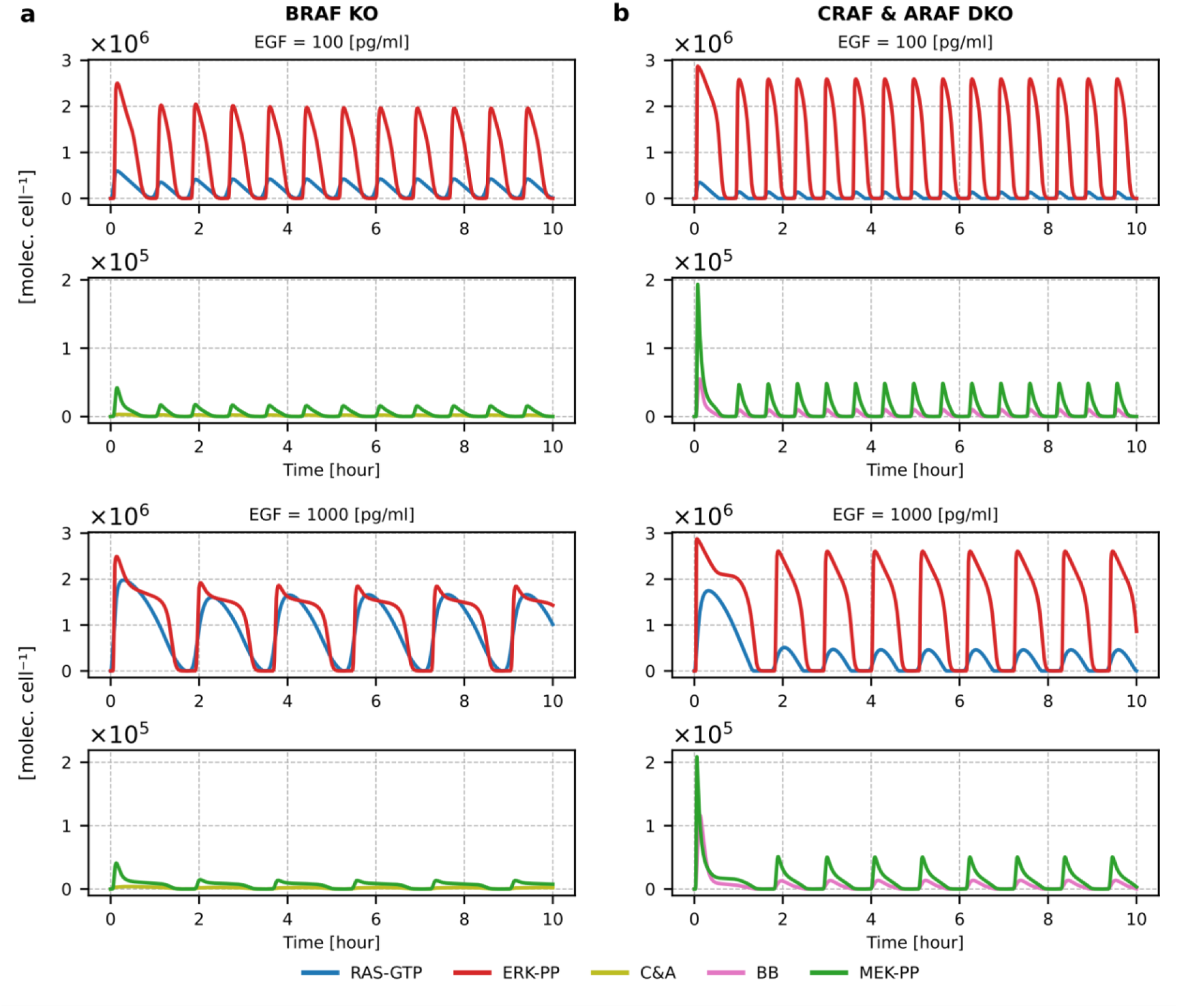
Relaxation oscillations in BRAF KO and ARAF & CRAF DKO cells. **(a)** Profiles of RAS-GTP, C&A dimers, MEKpp, and ERKpp in BRAF KO cells. **(b)** Profiles of RAS-GTP, BB dimers, MEKpp and ERKpp in ARAF & CRAF DKO cells. The corresponding plots for the low and high EGF concentrations, 20 pg/ml and 10000 pg/ml, are given in Fig. 5.

**Figure S3.**
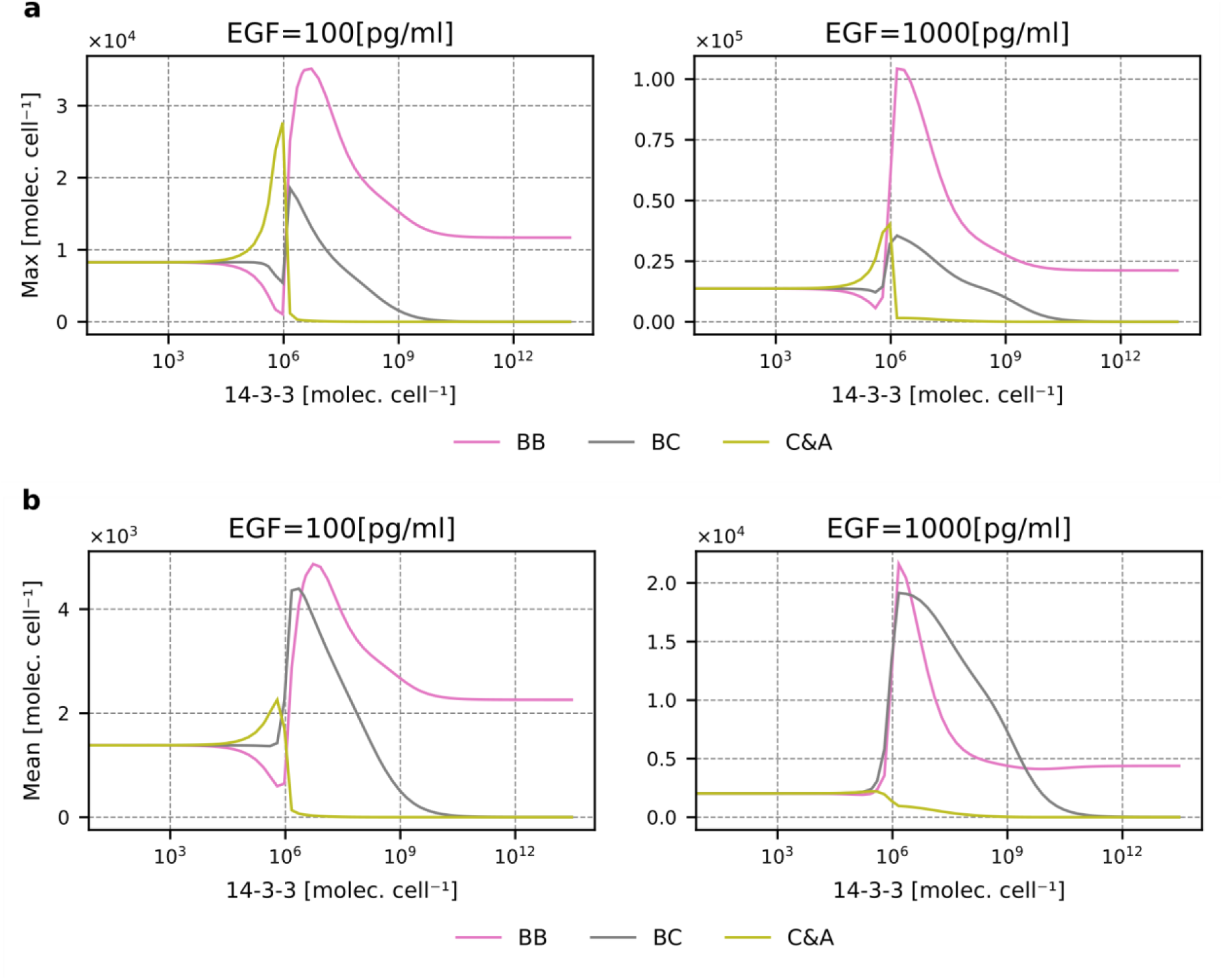
Abundance of RAF isoforms dimers as a function of 14-3-3 level. **(a)** The peak value of BB, BC and C&A dimers as a function of 14-3-3 for EGF concentrations 100, 1000 pg/ml. **(b)** The average value of BB, BC and C&A dimers (for the first 10 hours of EGF stimulation) as a function of 14-3-3 for EGF concentrations 100, 1000 pg/ml. The corresponding plots for the low and high EGF concentrations, 20 pg/ml and 10000 pg/ml, are given in Fig. 9.

**Figure S4.**
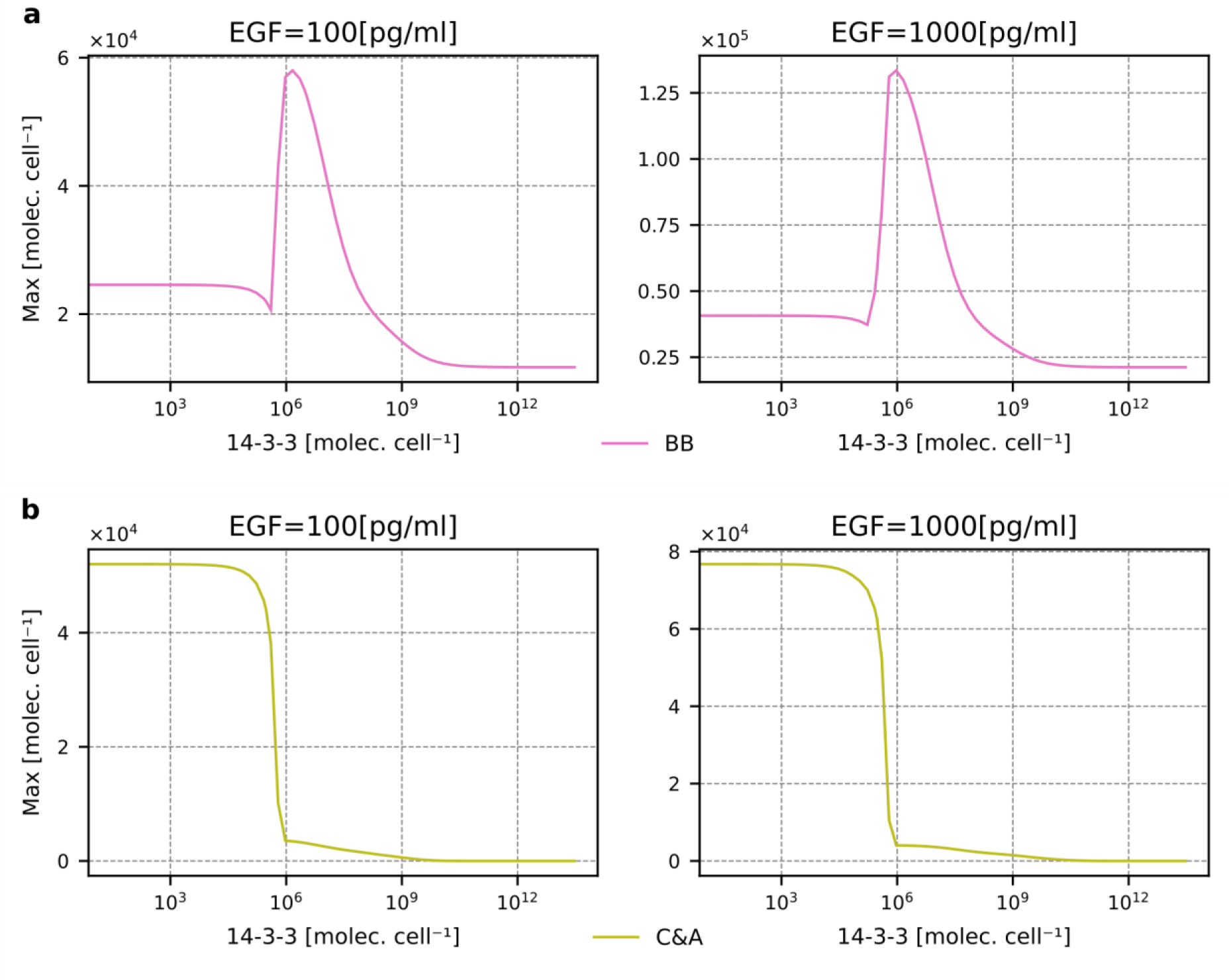
Abundance of RAF isoforms dimers as a function of 14-3-3 level for BRAF KO and CRAF & ARAF DKO cells. **(a)** The peak value of C&A dimers in BRAF KO cells as a function of 14-3-3 for EGF concentrations 100, 1000 pg/ml. **(b)** The peak value of BB dimers in CRAF & ARAF DKO cells as a function of 14-3-3 EGF concentrations 100, 1000 pg/ml. The corresponding plots for the low and high EGF concentrations, 20 pg/ml and 10000 pg/ml, are given in Fig. 1.

**Figure S5.**
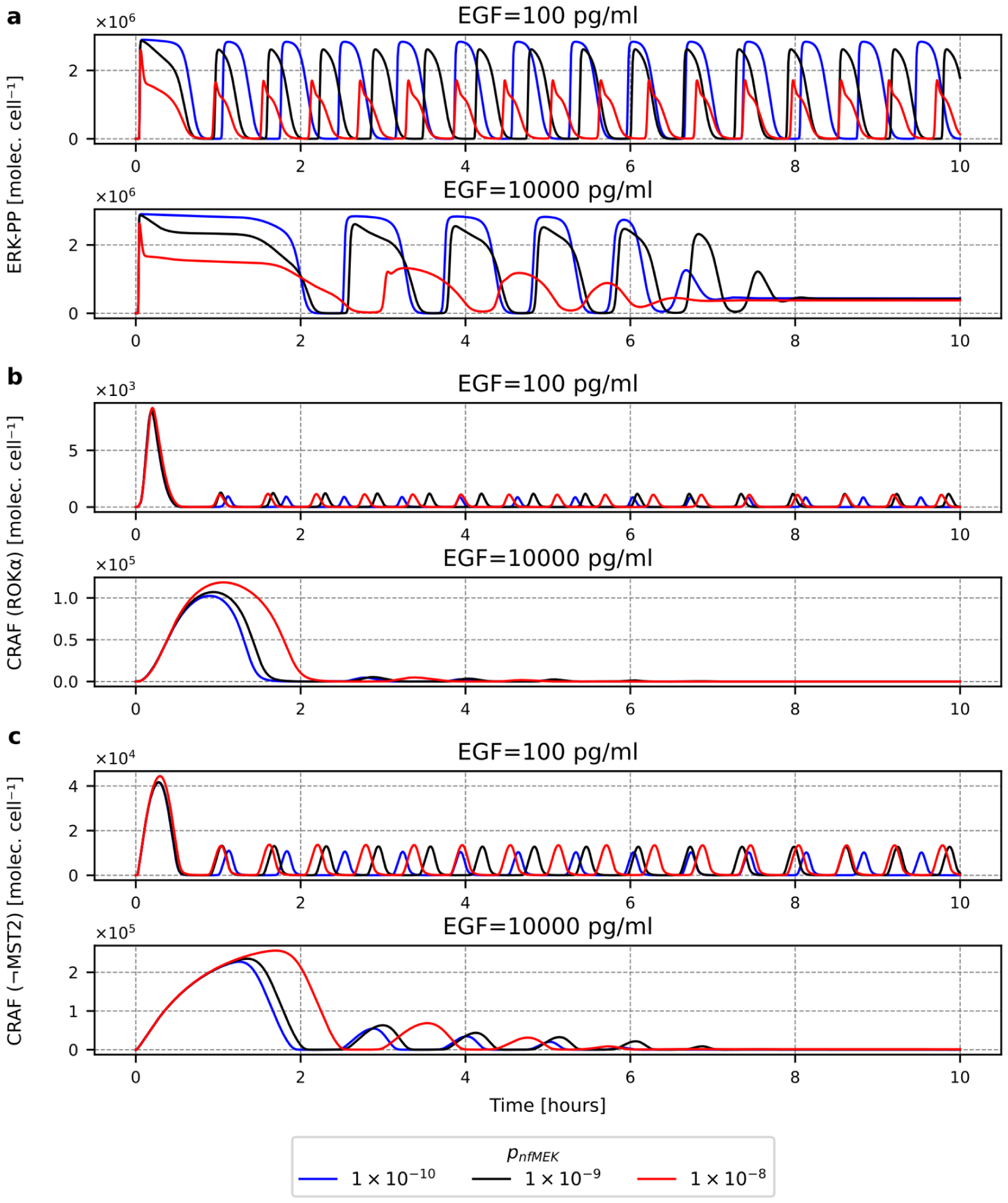
The impact of the ERK to MEK1 feedback strength on the time profiles of. **(a)** ERKpp, **(b)** CRAF in ROKα-compatible state, **(c)** CRAF in MST2-incompatible state.

